# YAP/TAZ activation in fibroblasts coordinates fibrotic remodeling, fibroinflammation, and epithelial dysfunction in pulmonary fibrosis

**DOI:** 10.1101/2025.06.23.661212

**Authors:** Masum M. Mia, Ashwatthaman Selvan, Uthayanan Nilanthi, Manvendra K. Singh

## Abstract

Idiopathic pulmonary fibrosis (IPF) is a fatal lung disease marked by progressive scarring with unknown causes and limited treatments. Myofibroblasts drive fibrosis by depositing excess matrix, however the mechanisms driving fibroblast-to-myofibroblast transformation and how myofibroblast-secreted factors disrupt the alveolar niche, undermining lung repair and regeneration remain poorly understood. Here we show that YAP and TAZ are activated in lung fibroblasts from pulmonary fibrosis patients and bleomycin-treated mice. Targeted deletion of *Yap/Taz* in fibroblasts significantly dampened the fibroinflammatory response, decreased myofibroblast activation, and ultimately resulted in attenuated fibrosis and enhanced regeneration of alveolar epithelial cells after bleomycin-induced injury. Conversely, fibroblast-specific overexpression of constitutively active YAP (YAP^5SA^) aggravated fibrosis by amplifying fibroinflammatory responses and simultaneously suppressing alveolar epithelial regeneration. Pharmacological inhibition of YAP/TAZ using verteporfin halted the development of bleomycin-induced fibrosis and even reversed established fibrosis in mice. Verteporfin effectively prevented fibroblast-to-myofibroblast transition and promoted collagen I degradation by lowering TIMP levels and enhancing activation of MMP1 and MMP9, demonstrating the significance of the YAP-TIMP-MMP1/9 axis in facilitating ECM breakdown. Furthermore, YAP activation in fibroblasts disrupted alveolar type II (AT2) cell homeostasis by inducing senescence through IGF1-IGF1R-mediated paracrine signaling. Blocking IGF1 signaling with a neutralizing antibody reduced the number of senescent AT2 cells, demonstrating the significance of the YAP-IGF1-IGF1R axis in maintaining alveolar epithelial cell homeostasis. Targeting YAP/TAZ may offer therapeutic strategies to mitigate pulmonary fibrosis by simultaneously mitigating pathological fibroblast activation, fibroinflammatory response, reducing AT2 cell senescence, and promoting alveolar epithelial regeneration.

## Introduction

Idiopathic pulmonary fibrosis (IPF) is a life-threatening chronic lung disease of unknown cause, marked by progressive scarring of the lung tissue. This fibrosis is caused by excessive extracellular matrix deposition between alveolar sacs, leading to destruction of peripheral lung tissue and impaired respiratory function (1). Complications associated with idiopathic pulmonary fibrosis (IPF) such as chronic hypoxia, fatigue, pain, coughing, and reduced mobility worsen over time, with advanced stages causing pulmonary hypertension and respiratory failure, further worsening prognosis (1–4). The median survival following a diagnosis of IPF is approximately 2 to 3 years, with higher mortality in older adults (4, 5). To date, two FDA-approved antifibrotic drugs, nintedanib and pirfenidone are available for the treatment of IPF (6). These drugs slow down IPF progression but cannot reverse established fibrosis (3, 7, 8). As IPF prevalence rises, the molecular and cellular mechanisms driving pulmonary fibrosis remain poorly understood, underscoring the urgent need for therapies that both halt progression and reverse fibrosis.

YAP and TAZ are transcriptional co-activators in the Hippo signaling pathway, acting as key sensors and integrators of mechanical and biochemical cues. Though initially characterized as master regulators of organ size during development, YAP and TAZ are now recognized as pivotal drivers of human disease pathologies. Beyond their developmental roles, these transcriptional co-activators fuel cancer progression, impair tissue regeneration, and critically orchestrate fibrotic disorders across organs. In injury-induced fibrotic conditions, intercellular crosstalk critically shapes the fibroinflammatory response, ultimately determining disease progression. Dynamic interactions between fibroblasts, epithelial cells and immune cells (e.g., macrophages), create a self-amplifying loop of inflammation and fibrosis. This dysregulated crosstalk not only drives persistent inflammation and pathological remodeling but also impairs resolution mechanisms, making it a pivotal determinant of fibrosis progression across organs, including pulmonary fibrosis. Targeting these cell-specific interactions may offer novel therapeutic strategies to disrupt the fibroinflammatory cascade. A recent study demonstrated that activation of YAP/TAZ in alveolar type 2 (AT2) epithelial cells is indispensable for initiating proper alveolar repair in response to bleomycin-induced injury (9). Genetic deletion of *Yap/Taz* in AT2 cells leads to pathological alveolar remodeling following bleomycin injury, characterized by impaired AT2-to-AT1 cell differentiation, increased collagen deposition, exaggerated maladaptive fibroinflammatory responses, and higher mortality (9). These findings demonstrate that AT2-specific YAP/TAZ activity is crucial for proper epithelial regeneration. Recently we demonstrated that genetic deletion of YAP/TAZ in myeloid cells conferred significant protection against bleomycin-induced pulmonary fibrosis through coordinated multi-tiered effects (10). The genetic ablation disrupted monocyte-derived alveolar macrophage (Mo-AM) recruitment to injured lungs while simultaneously impairing their capacity to mount robust inflammatory responses through attenuated cytokine/chemokine production. This myeloid-specific perturbation created a tissue microenvironment more conducive to regeneration, as evidenced by enhanced alveolar epithelial repair (10). While YAP/TAZ are established mediators of fibrosis across multiple organs, their fibroblast-specific contributions to pulmonary fibrosis remain poorly characterized. Myofibroblasts, the primary effector cells of fibrosis, drive disease progression through pathological extracellular matrix deposition. However, the mechanisms by which YAP/TAZ orchestrate fibroblast-to-myofibroblast transformation and how myofibroblast-secreted factors subsequently disrupt the alveolar niche to undermining lung repair and regeneration are still undefined.

Our present study elucidates the critical role of YAP/TAZ in lung fibroblasts during bleomycin-induced pulmonary fibrosis (PF). Fibroblast-specific deletion of Yap/Taz attenuated fibrosis by reducing macrophage infiltration, suppressing inflammatory cytokine/chemokine expression, and diminishing profibrotic myofibroblast activation and extracellular matrix (ECM) accumulation, while simultaneously promoting alveolar epithelial regeneration. Whereas constitutive YAP activation (YAP^5SA^) exacerbates PF by amplifying macrophage recruitment, inflammatory cytokine/chemokine expression, profibrotic myofibroblast activation and impairing alveolar regeneration. We found that YAP activation in fibroblasts disrupts alveolar regeneration by suppressing AT2 alveolar epithelial cell proliferation while promoting cellular senescence through IGF1-IGF1R signaling. Pharmacological inhibition of YAP/TAZ with verteporfin demonstrated dual therapeutic efficacy by both preventing the onset and reversing established lung fibrosis. Verteporfin suppressed myofibroblast activation, promoted collagen degradation through modulation of the MMP1/MMP9-TIMP axis, attenuated pro-inflammatory transcription factor signaling, and restored alveolar epithelial cell populations. Collectively, these findings demonstrate that YAP/TAZ play a central role in regulating fibroinflammatory response and myofibroblast-epithelial crosstalk in pulmonary fibrosis, orchestrating both disease progression and resolution. Our findings not only elucidate the pathogenic role of YAP/TAZ in maintaining the fibrotic niche but also validate their therapeutic potential as druggable targets for clinical intervention in fibrotic lung diseases.

## Results

### YAP/TAZ are activated in lung fibroblasts from patients with pulmonary fibrosis and in bleomycin-treated mice

To determine the role of YAP/TAZ in fibrotic progression, we first compared their activation status in fibroblasts from healthy and fibrotic human lungs. Immunofluorescence analysis demonstrated significantly enhanced nuclear accumulation of both YAP and TAZ in PDGFRα+ fibroblasts from fibrotic lungs relative to healthy controls (Figure 1A-C). This activation pattern was further validated in DDR2+Vimentin+ fibroblasts, which exhibited markedly increased nuclear YAP/TAZ expression in fibrotic samples (Supplementary Figure 1A).

**Figure 1:**
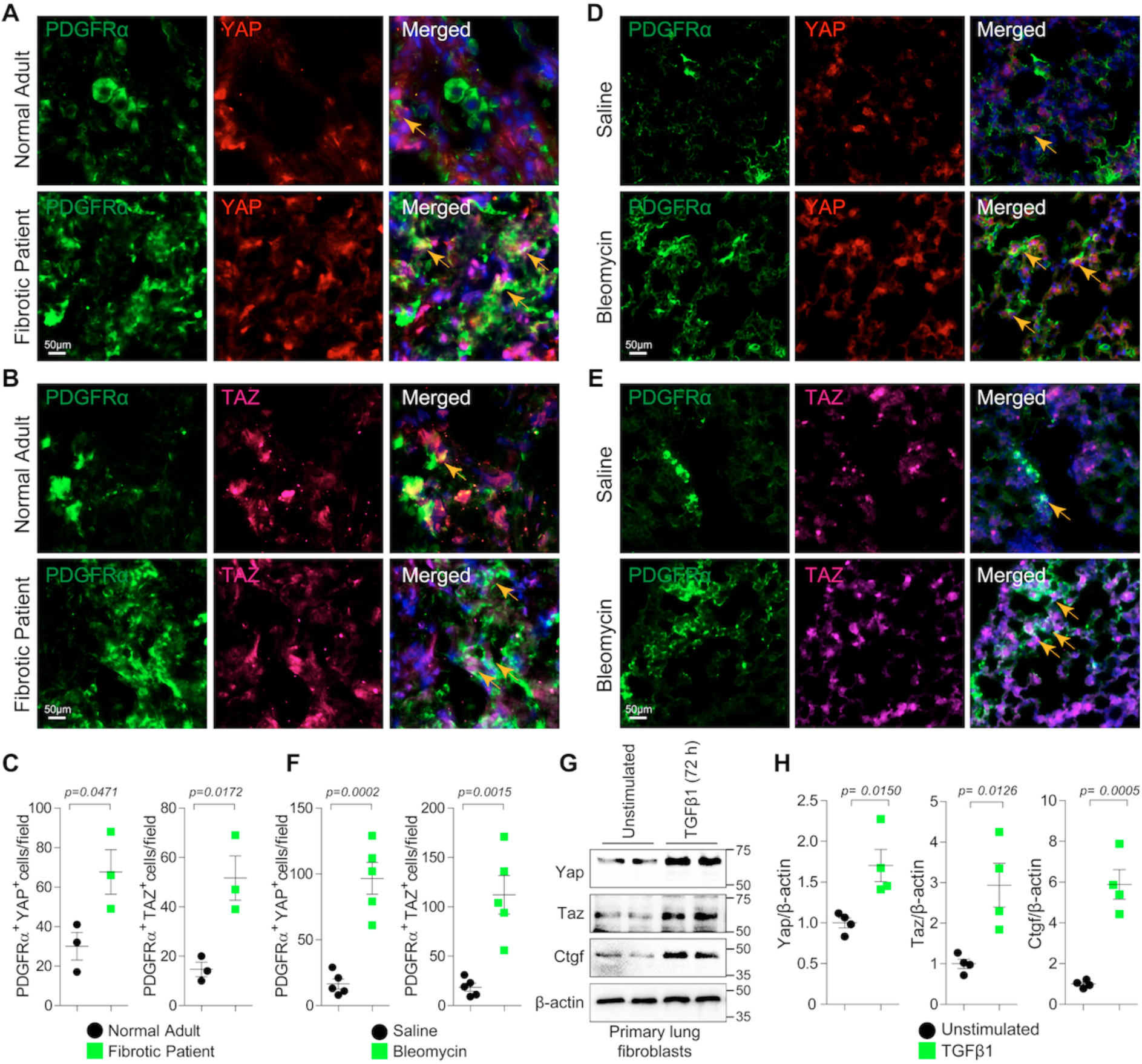
YAP/TAZ are activated in lung Fibroblasts. (A-C) Immunofluorescence staining and quantification of YAP or TAZ co-stained with PDGFRα in lung sections from human patients diagnosed with pulmonary fibrosis compared with normal adult human lung (n = 3). (D-F) Immunofluorescence staining and quantification of YAP or TAZ co-stained with PDGFRα in wildtype mice lung sections at 7 days post-bleomycin injury compared to saline-treated controls (n = 5). (G-H) Immunoblot analysis of YAP, TAZ, and CTGF on primary mouse lung fibroblasts treated with/without TGFβ1 (10 ng/mL) for 72 hours (n = 4). Quantification of indicated proteins relative to β-actin. The data are represented as the mean ± SEM; comparison by two-tailed unpaired t-test. *, *p* < 0.05; **, *p* < 0.01; ***, *p* < 0.001; NS, not significant.

To define the fibroblast-specific functions of YAP/TAZ in pulmonary fibrosis, we analyzed their activation in a bleomycin-induced mouse model. Immunohistochemistry of lung sections at 7 days post-injury revealed significant nuclear accumulation of YAP/TAZ in PDGFRα+ fibroblasts (Figure 1D-F) and DDR2+Vimentin+ fibroblasts (Supplementary Figure 1B) compared to saline-treated controls. To further support this finding, we treated primary mouse lung fibroblasts with TGFβ1, a canonical driver of fibrogenesis. Immunoblot analysis confirmed upregulation of both YAP and TAZ proteins, along with their established downstream target CTGF (Figure 1G-H), confirming activation of this signaling axis in response to profibrotic stimulation. Together, these findings demonstrate that YAP/TAZ are robustly activated in lung fibroblasts under profibrotic conditions and likely modulate their phenotypic transformation.

### Genetic inactivation of YAP/TAZ attenuates the inflammatory response induced by bleomycin in the lung

To investigate YAP/TAZ function in lung fibroblasts, we generated tamoxifen-inducible, fibroblast-specific Yap/Taz double-knockout mice (*Col1a2^Cre(ER)T^;Yap^flox/flox^; Taz^flox/flox^*) by crossing floxed Yap/Taz mice (*Yap^flox/flox^;Taz^flox/flox^*) with *Col1a2^Cre(ER)T^* mice (Figure 2A). Following saline or bleomycin administration (day 0), mice received daily tamoxifen injections (days 1-8) with tissue collection 6 days post-final injection. Immunofluorescence analysis of bleomycin-treated lung sections revealed reduced nuclear expression of YAP and TAZ in PDGFRα+ fibroblasts of *Col1a2^Cre(ER)T^; Yap^flox/flox^;Taz^flox/flox^* mice compared to controls, indicating efficient deletion of YAP/TAZ in fibroblasts (Supplementary Figure 2A–B).

**Figure 2:**
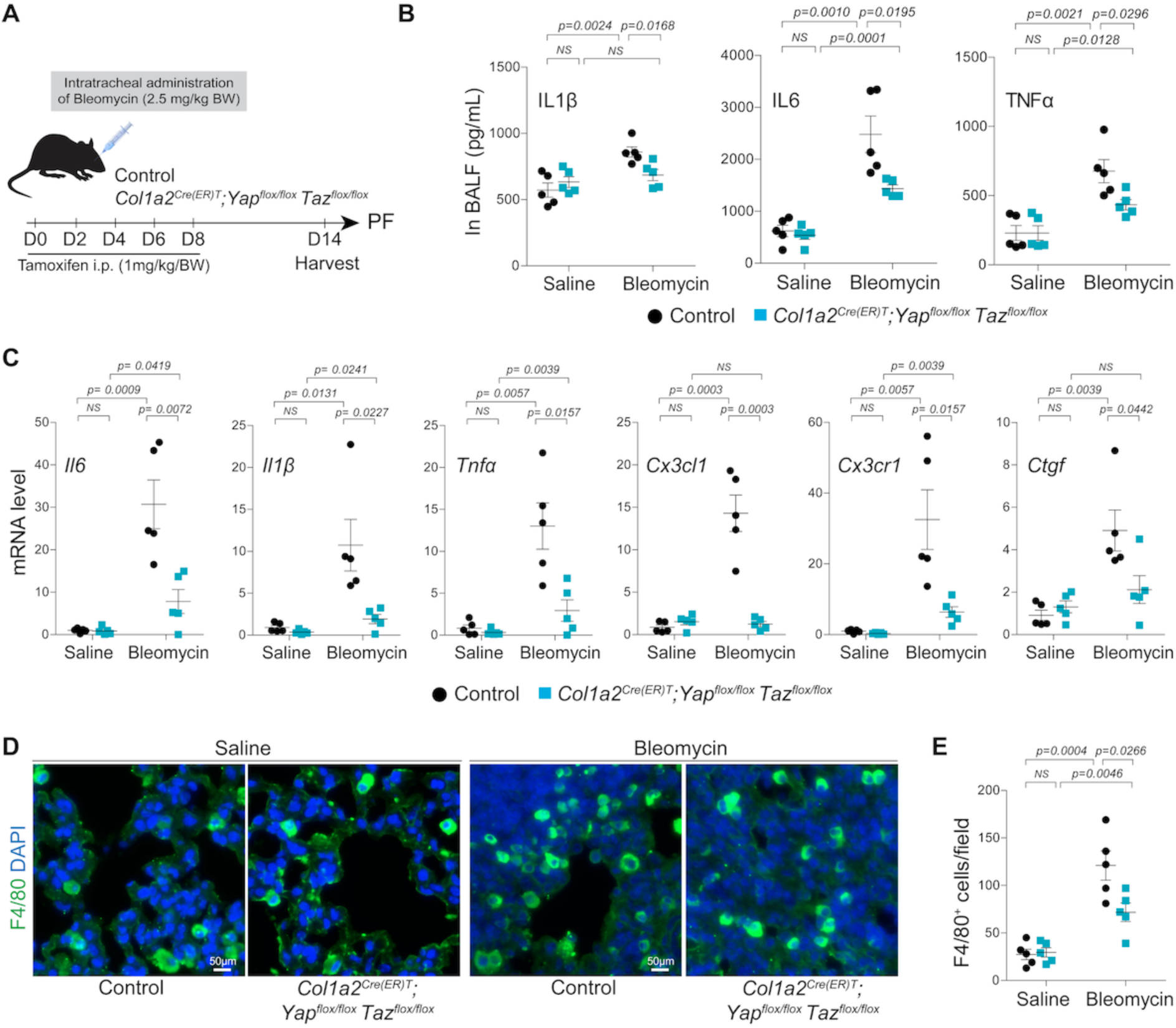
*Yap/Taz* inactivation in lung fibroblasts impairs bleomycin-induced inflammatory response in mice. (A) Graphical presentation of experimental design illustrating the bleomycin induced lung fibrosis; Both *Yap^flox/flox^; Taz^flox/flox^* (control) and *Col1a2^Cre(ER)T^; Yap^flox/flox^; Taz^flox/flox^* (*dKO*) mice were challenged to one dose of intratracheal administration of bleomycin (2.5 mg/kg body weight) or saline with following intraperitoneal injection of tamoxifen (1 mg per kg body weight/day) as indicated time points. Lung tissue and bronchoalveolar lavage fluid (BALF) were harvested at 14 days post-bleomycin injury for further analysis. (B) Enzyme-linked immunosorbent assay (ELISA) to measure the protein release of pro-inflammatory cytokines IL1β, IL6 and TNF-α in BALF collected from control and *dKO* mice (n = 5) treated with saline or bleomycin for 14 days. (C) Real-time qPCR for pro-inflammatory cytokines and chemokines such as *Il6*, *Il1β*, *Tnfα, Cx3cl1,* and *Cx3cr1*; with Yap/Taz target gene *Ctgf* using lung tissue RNA isolated from control and *dKO* mice (n = 5) treated with saline or bleomycin for 14 days. (D-E) Immunostaining and quantification of F4/80 positive macrophage infiltration on the lung from control and *dKO* mice (n = 5) treated with saline or bleomycin for 14 days. The data are represented as the mean ± SEM; comparison by two-tailed unpaired t-test. *, *p* < 0.05; **, *p* < 0.01; ***, *p* < 0.001; NS, not significant.

To assess the impact of fibroblast-specific YAP/TAZ inactivation on bleomycin-induced inflammation, we treated control and *Col1a2^Cre(ER)T^;Yap^flox/flox^;Taz^flox/flox^*mice with bleomycin or saline for 14 days. Analysis of bronchoalveolar lavage fluid (BALF) by ELISA revealed comparable levels of IL-1β, IL-6, and TNF-α between control and knockout mice under saline conditions (Figure 2B). However, bleomycin-challenged Yap/Taz-deficient mice showed significantly reduced BALF levels of these cytokines compared to controls (Figure 2B). To further characterize the inflammatory profile, we performed qRT-PCR on lung tissue. Expression of key inflammatory cytokines and chemokines including *Il6, Il1β, Tnfα, Cx3cl1*, and *Cx3cr1* was significantly downregulated in YAP/TAZ-deficient lungs compared to controls. Additionally, expression of the YAP/TAZ target gene *Ctgf* was markedly reduced (Figure 2C), confirming effective gene inactivation and functional suppression of YAP/TAZ signaling. We next assessed macrophage infiltration by immunofluorescence staining for F4/80. While saline-treated groups showed no differences, bleomycin treatment led to increased F4/80⁺ macrophage accumulation in control lungs, which was significantly attenuated in YAP/TAZ-deficient mice (Figure 2D-E). Collectively, these results establish YAP/TAZ as critical regulators of fibroinflammatory responses in lung injury, modulating both cytokine production and immune cell recruitment.

### Genetic inactivation of YAP/TAZ attenuates bleomycin-induced lung fibrosis

To evaluate the impact of fibroblast-specific YAP/TAZ deletion on lung fibrosis, we conducted histological, immunohistochemical, immunoblotting, and qRT-PCR analyses on lung tissues from control and *Col1a2^Cre(ER)T^;Yap^flox/flox^;Taz^flox/flox^*mice at 14 days post-bleomycin instillation. Fibrosis severity was markedly reduced in YAP/TAZ-deficient mice, as evidenced by diminished Sirius Red staining and significantly lower Ashcroft scores compared to control mice (Figure 3A-B). To quantitatively evaluate collagen deposition, we performed a hydroxyproline assay on lung tissue. Bleomycin administration led to a robust increase in hydroxyproline content in control lungs, reflecting excessive collagen accumulation. In contrast, YAP/TAZ-deficient lungs showed a significant reduction in hydroxyproline levels at day 14 post-injury (Figure 3C), indicating attenuated fibrotic remodeling. Gene expression analyses showed that bleomycin markedly increased the mRNA levels of extracellular matrix (ECM)-related genes such as *Col1a1* (collagen type I) and *Periostin*, as well as myofibroblast-associated genes including *Acta2* (α-smooth muscle actin) and *Tagln* (SM22α) in control lungs compared to saline-treated mice. In contrast, expression of these genes was significantly reduced in YAP/TAZ-deficient lungs at 14 days post-bleomycin injury. No significant differences in gene expression were observed between saline-treated control and YAP/TAZ-deficient lungs (Figure 3D). To validate these findings further, we assessed expression of collagen I and αSMA by immunofluorescence and immunoblotting. Bleomycin exposure led to a pronounced increase in both proteins in control lungs, whereas their expression was significantly attenuated in YAP/TAZ-deficient lungs. No differences were detected between saline-treated groups (Figure 3E-H). These results suggest that YAP/TAZ-dependent pro-fibrotic activity in lung fibroblasts promotes fibrosis after injury.

**Figure 3:**
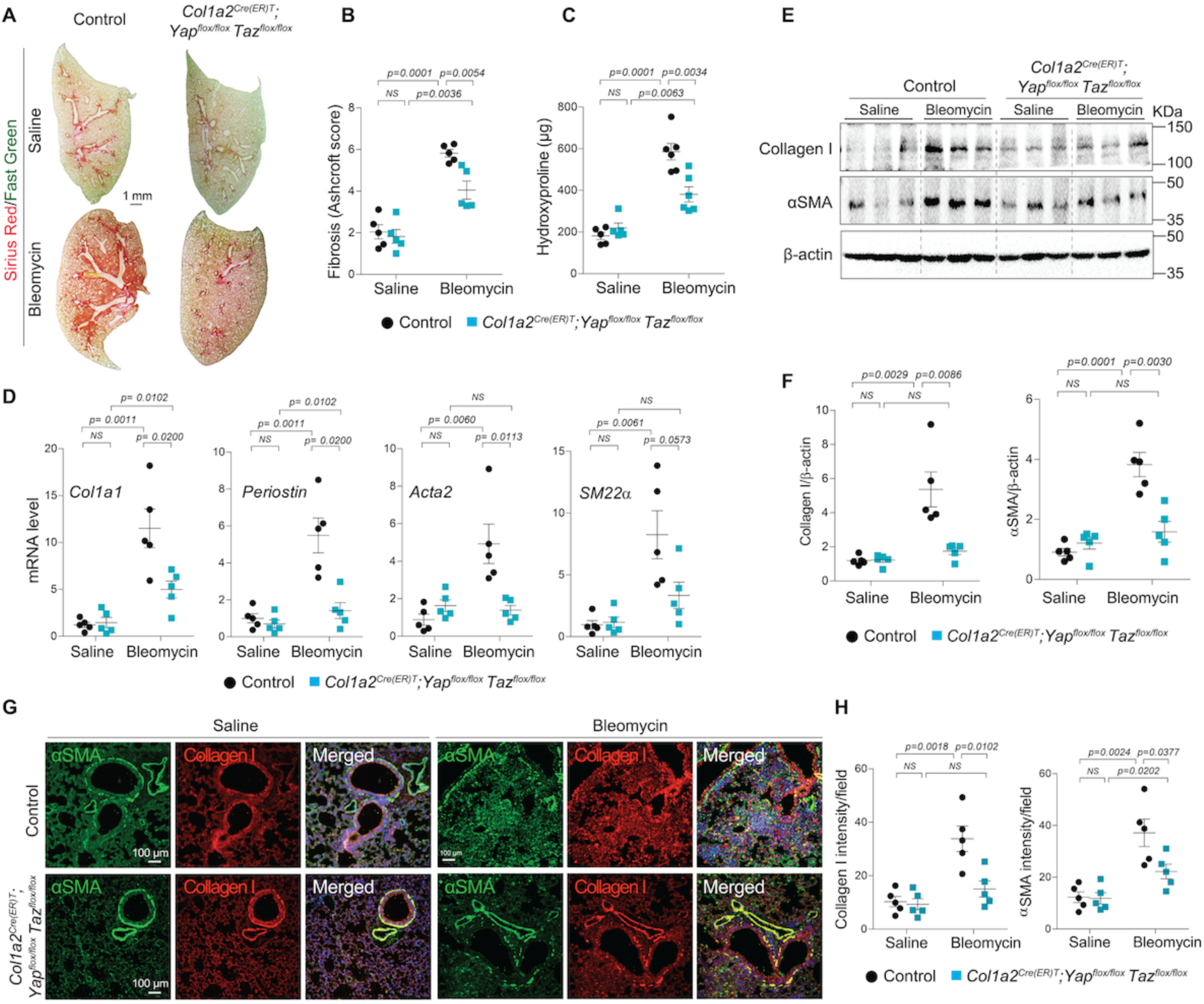
Fibroblasts-specific *Yap/Taz* inactivation attenuates bleomycin-induced lung fibrosis. (A-B) Sirius Red/Fast Green staining and quantification (presented as Ashcroft score) on lung sections from control and *dKO* mice (n = 5) treated with saline or bleomycin for 14 days. (C) Quantification of hydroxyproline content using lung tissue from control and *dKO* mice (n = 5-6) treated with saline or bleomycin for 14 days. (D) Real-time qPCR for profibrotic genes *Acta2, SM22α, Col1a1,* and *Periostin* on lung tissue from control and *dKO* mice (n = 5) treated with saline or bleomycin for 14 days. (E-F) Immunoblot analysis and quantification of αSMA and collagen I protein using lung tissue lysate from control and *dKO* mice (n = 5) treated with saline or bleomycin for 14 days. Quantification of indicated proteins relative to β-actin. (G-H) Immunofluorescence staining and quantification of αSMA and collagen I in lung sections from control and *dKO* mice (n = 5) treated with saline or bleomycin for 14 days. Fluorescence intensity was quantified with imageJ software. The data are represented as the mean ± SEM; comparison by two-tailed unpaired t-test. *, *p* < 0.05; **, *p* < 0.01; ***, *p* < 0.001; NS, not significant.

### Fibroblast-specific overexpression of YAP exacerbates bleomycin-induced lung inflammation and fibrosis

To investigate the role of YAP in lung fibrosis, we selectively expressed a constitutively active form of YAP in fibroblasts by crossing *Col1a2^Cre(ER)T^* mice with *R26^YAP5SA^* mice, generating *Col1a2^Cre(ER)T^;R26^YAP5SA^*mice. Immunostaining of control (*R26^YAP5SA^*) and *Col1a2^Cre(ER)T^;R26^YAP5SA^*lung sections confirmed robust nuclear YAP localization in PDGFRα+ lung fibroblasts (Supplementary Figure 3), confirming successful fibroblast-specific YAP activation. To assess the impact of fibroblast YAP activation on inflammation and fibrosis, mice were treated with saline or bleomycin on day 0, followed by tamoxifen injections until day 8. Lungs were harvested and analyzed six days after the last tamoxifen dose (Figure 4A). qRT-PCR analysis revealed that YAP overexpression significantly upregulated mRNA expression of pro-inflammatory cytokines (*Il1β, Il6, TNF-α*), chemokines (*Cx3cl1, Cx3cr1*), and the canonical YAP/TAZ target gene *Ctgf* under both saline and bleomycin conditions (Figure 4B). We next evaluated macrophage infiltration by immunostaining for F4/80. Compared to controls, *Col1a2^Cre(ER)T^;R26^YAP5SA^*lungs exhibited increased F4/80+ macrophages after both saline and bleomycin treatment (Figure 4C-D), indicating that fibroblast YAP activation promotes macrophage recruitment.

**Figure 4:**
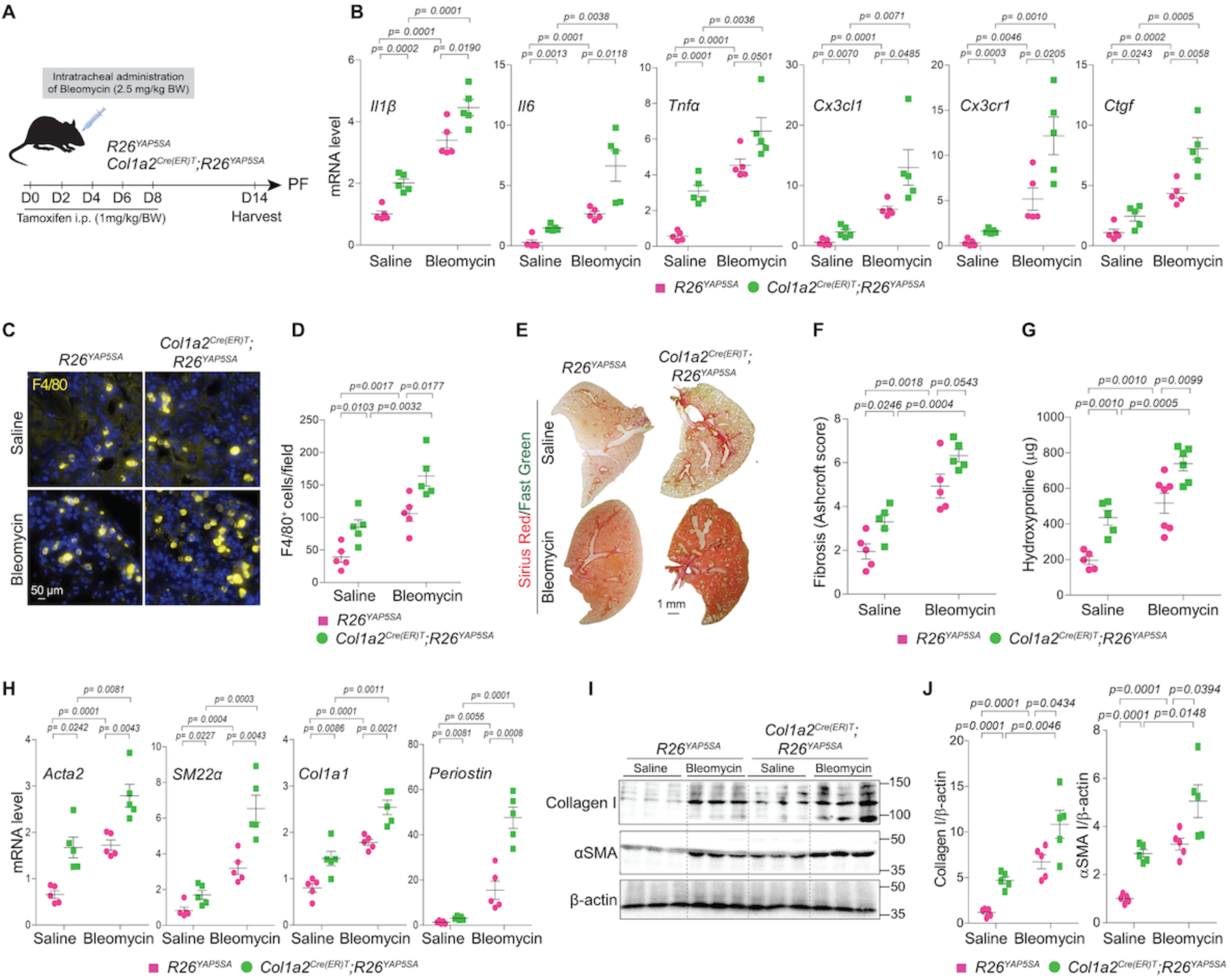
*Yap* overexpression in fibroblasts exacerbates bleomycin-induced lung inflammation and fibrosis in mice. (A) Graphical presentation of experimental design illustrating the bleomycin induced lung fibrosis; Both *R26^YAP5SA^* (control) and *Col1a2^Cre(ER)T^; R26^YAP5SA^* (Yap over expressing mice) were challenged to one dose of intratracheal administration of bleomycin (2.5 mg/kg body weight) or saline with following intraperitoneal injection of tamoxifen (1 mg per kg body weight/day) as indicated time points. Lungs were harvested at 14 days post-bleomycin injury for further analysis. (B) Real-time qPCR for pro-inflammatory cytokines and chemokines such as *Il6*, *Il1β*, *Tnfα, Cx3cl1,* and *Cx3cr1*; with Yap/Taz target gene *Ctgf* using lung tissue RNA isolated from *R26^YAP5SA^* and *Col1a2^Cre(ER)T^; R26^YAP5SA^*mice (n = 5) treated with saline or bleomycin for 14 days. (C-D) Immunostaining and quantification of F4/80 positive macrophage infiltration on the lung from *R26^YAP5SA^* and *Col1a2^Cre(ER)T^; R26^YAP5SA^*mice (n = 5) treated with saline or bleomycin for 14 days. (E-F) Sirius Red/Fast Green staining and quantification (presented as Ashcroft score) on lung sections from *R26^YAP5SA^* and *Col1a2^Cre(ER)T^; R26^YAP5SA^* mice (n = 5) treated with saline or bleomycin for 14 days. (G) Quantification of hydroxyproline content using lung tissue from *R26^YAP5SA^* and *Col1a2^Cre(ER)T^; R26^YAP5SA^* mice (n = 5-6) treated with saline or bleomycin for 14 days. (H) Real-time qPCR for profibrotic genes *Acta2, SM22α, Col1a1,* and *Periostin* on lung tissue from *R26^YAP5SA^* and *Col1a2^Cre(ER)T^; R26^YAP5SA^* mice (n = 5) treated with saline or bleomycin for 14 days. (I-J) Immunoblot analysis and quantification of collagen I and αSMA protein levels using lung tissue lysate from *R26^YAP5SA^* and *Col1a2^Cre(ER)T^; R26^YAP5SA^* mice (n = 5) treated with saline or bleomycin for 14 days. Quantification of indicated proteins relative to β-actin. The data are represented as the mean ± SEM; comparison by two-tailed unpaired t-test. *, *p* < 0.05; **, *p* < 0.01; ***, *p* < 0.001; NS, not significant.

Next, we assessed the extent of fibrosis in response to fibroblast-specific YAP overexpression. Histological evaluation revealed markedly exacerbated lung fibrosis in *Col1a2^Cre(ER)T^;R26^YAP5SA^*mice compared to controls, as evidenced by increased Sirius Red staining and significantly elevated Ashcroft scores, both at baseline and 14 days post-bleomycin administration (Figure 4E–F). Consistent with these findings, hydroxyproline assays revealed elevated collagen content in *Col1a2^Cre(ER)T^;R26^YAP5SA^*lungs under both saline and bleomycin-treated conditions (Figure 4G). To further delineate the molecular basis of this profibrotic effects of YAP, we examined the expression of key fibrogenic genes. YAP overexpression significantly increased mRNA levels of *Acta2*, *SM22α*, *Col1a1*, and *Periostin* (Figure 4H). Consistent with transcriptomic findings, immunoblot analysis revealed elevated levels of Collagen I and α-SMA in *Col1a2^Cre(ER)T^;R26^YAP5SA^*lungs (Figure 4I–J). Together, these results demonstrate that fibroblast-specific YAP activation drives fibro-inflammatory responses, enhances macrophage infiltration, and amplifies profibrotic fibroblast activity, underscoring its critical role in lung fibrosis pathogenesis.

### YAP/TAZ activity in alveolar fibroblasts regulates alveolar type 2 (AT2) epithelial cell proliferation and senescence

To evaluate the impact of fibroblast-specific Yap/Taz deletion on alveolar epithelial cell behaviour following lung injury, we analyzed lung tissues from control and *Col1a2^Cre(ER)T^;Yap^flox/flox^;Taz^flox/flox^*mice 14 days after bleomycin exposure. Immunohistochemical evaluation revealed substantial epithelial damage following injury, characterized by a marked reduction in both AT1 (Hopx+) and AT2 (pro-SPC+) cells compared to saline-treated controls (Figure 5A). We observed a significant loss of AT2 cells post-injury in both groups; however, Yap/Taz-deficient lungs exhibited a higher number of AT2 cells relative to controls. Similarly, AT1 cell numbers were also higher in Yap/Taz-deficient lungs compared to injured controls. These findings suggest that fibroblast-specific Yap/Taz deletion promotes epithelial cell preservation or regeneration and may contribute to improved alveolar repair following fibrotic lung injury.

**Figure 5:**
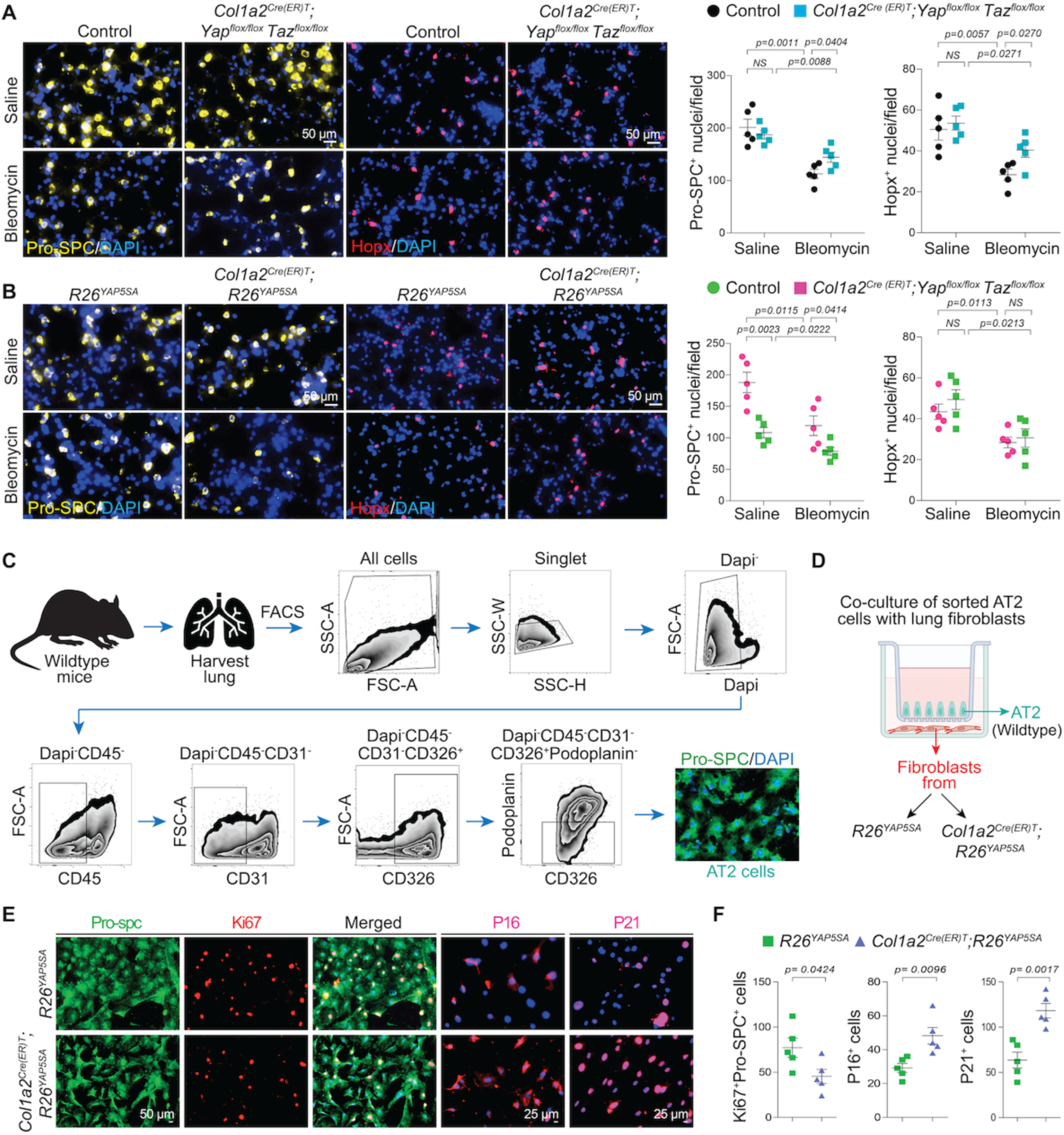
Modulation of Yap/Taz activity in lung fibroblasts affects alveolar epithelial cell proliferation and senescence. (A) Immunostaining and quantification of pro-surfactant protein C (Pro-SPC)+ AT2-cells and Hopx+ AT1-cells in control *versus dKO* mouse lung (n=5) treated with saline or bleomycin for 14 days. (B) Immunostaining and quantification of Pro-SPC+ AT2-cells and Hopx+ AT1-cells in *R26^YAP5SA^ versus Col1a2^Cre(ER)T^; R26^YAP5SA^* mouse lung (n=5) treated with saline or bleomycin for 14 days. (C-D) Graphical presentation for flow cytometry gating strategy to isolate AT2 cells from wild-type mice lungs defined by Dapi−CD45−CD31−CD326+Podoplanin− and subsequently co-cultured with lung fibroblasts from *R26^YAP5SA^* or *Col1a2^Cre(ER)T^; R26^YAP5SA^* lungs. (E-F) Immunostaining and quantification of Pro-SPC+Ki67+, or p16+ and p21+ AT2-cells from lungs on co-culture insert where fibroblasts were cultured underneath of sorted AT2 cells (n=5) for 14 days. Data are presented as mean±SEM, with statistical comparisons made using a two-tailed unpaired t-test. Significance levels are indicated with p-value (significant at p<0.05).

To further determine whether YAP overexpression in fibroblasts influences alveolar epithelial cell behaviour, control and *Col1a2^Cre(ER)T^;R26^YAP5SA^*mice were treated with either saline or bleomycin for 14 days. Lungs were harvested and analyzed using immunohistochemistry, flow cytometry, and fibroblast-AT2 cell co-culture experiments. Immunostaining revealed a substantial loss of AT2 cells in response to bleomycin injury. Notably, YAP overexpression in fibroblasts exacerbated this loss, with a significant reduction in AT2 cells observed in both saline- and bleomycin-treated *Col1a2^Cre(ER)T^;R26^YAP5SA^*lungs compared to controls (Figure 5B). Similarly, the number of AT1 cells decreased following bleomycin injury; however, no significant difference in AT1 cell numbers was detected between control and *Col1a2^Cre(ER)T^;R26^YAP5SA^*mice in either treatment group (Figure 5B). Together, these findings underscore a detrimental role for fibroblast-specific YAP activation in impairing epithelial regeneration following lung injury. To further establish that the reduction in AT2 cell numbers was directly attributable to YAP activation in alveolar fibroblasts, we conducted fibroblast-AT2 cell co-culture experiments. Wild-type AT2 cells (defined by the surface markers DAPI⁻CD45⁻CD31⁻CD326⁺Podoplanin⁻) were isolated from mouse lungs via flow cytometry and co-cultured with lung fibroblasts isolated from *R26^YAP5SA^*and *Col1a2^Cre(ER)T^;R26^YAP5SA^* mice (Figure 5C-D). Following the co-culture period, AT2 cells were assessed by immunostaining for the proliferation marker Ki67 and the senescence markers p16 (Cdkn2a) and p21 (Cdkn1a) (Figure 5E). Co-immunostaining for pro-SPC and Ki67 revealed that co-culture with YAP-overexpressing fibroblasts significantly reduced the number of proliferating AT2 cells (pro-SPC⁺Ki67⁺). In contrast, YAP-overexpressing fibroblasts markedly increased the expression of senescence markers p21 and p16 in AT2 cells, compared to the controls (Figure 5E-F). These findings suggest that the secretome of YAP-activated fibroblasts suppresses AT2 cell proliferation while promoting their senescence.

### Increased fibroblast-specific YAP activity affects AT2 cell homeostasis through IGF1-IGF1R-mediated paracrine signaling

To elucidate the mechanisms underlying YAP-mediated fibroblast–epithelial crosstalk, we performed a candidate-based screen of fibroblast-derived factors and identified insulin-like growth factor 1 (IGF1) as a key target that may modulate alveolar epithelial cell behavior in a YAP-dependent manner. To investigate whether YAP regulates IGF1-IGF1R signaling during pulmonary fibrosis, we examined lung tissue samples from saline and bleomycin-treated mice. Immunoblot analysis for YAP, IGF1, and IGF1R revealed that bleomycin-induced lung injury markedly increased protein expression of IGF1 and IGF1R, along with YAP (Figure 6A). Primary lung fibroblasts from bleomycin-treated mice showed elevated mRNA levels of fibrotic markers such as *Acta2*, *Col1a1* and *Igf1* compared to saline controls (Figure 6B). Furthermore, using YAP-overexpressing fibroblasts, we confirmed YAP-dependent IGF1 upregulation at the protein level (Figure 6C). Immunofluorescence analysis revealed increased PDGFRα+IGF1+ fibroblasts and Pro-SPC+IGF1R+ alveolar epithelial cells following bleomycin injury (Figure 6D, Supplementary Figure 4A-B), establishing a potential paracrine signaling axis. Based on elevated IGF1 expression in YAP-overexpressing fibroblasts (Figure 6C), we postulated that YAP drives fibroblast-mediated AT2 senescence via IGF1-IGF1R signaling. Supporting this hypothesis, recombinant IGF1 treatment suppressed AT2 cell proliferation and induced senescence, as evidenced by increased populations of P21⁺ and P16⁺ AT2 cells (Figures 6E–F). To further investigate this mechanism, we co-cultured sorted AT2 cells (DAPI⁻CD45⁻CD31⁻CD326⁺Podoplanin⁻) with *Col1a2^Cre(ER)T^;R26^YAP5SA^*fibroblasts in the presence of either an IGF1-neutralizing antibody or an IgG control (Figures 6G–H). IGF1 neutralization significantly reduced the number of P21⁺ and P16⁺ AT2 cells induced by the YAP-overexpressing fibroblast secretome, while increasing the number of proliferating Pro-SPC⁺Ki67⁺ AT2 cells compared to IgG control treatment (Figures 6I–J). These findings suggest that YAP activation in fibroblasts effects AT2 cell proliferation and senescence via the IGF1-IGF1R signaling axis during bleomycin-induced pulmonary fibrosis.

**Figure 6.**
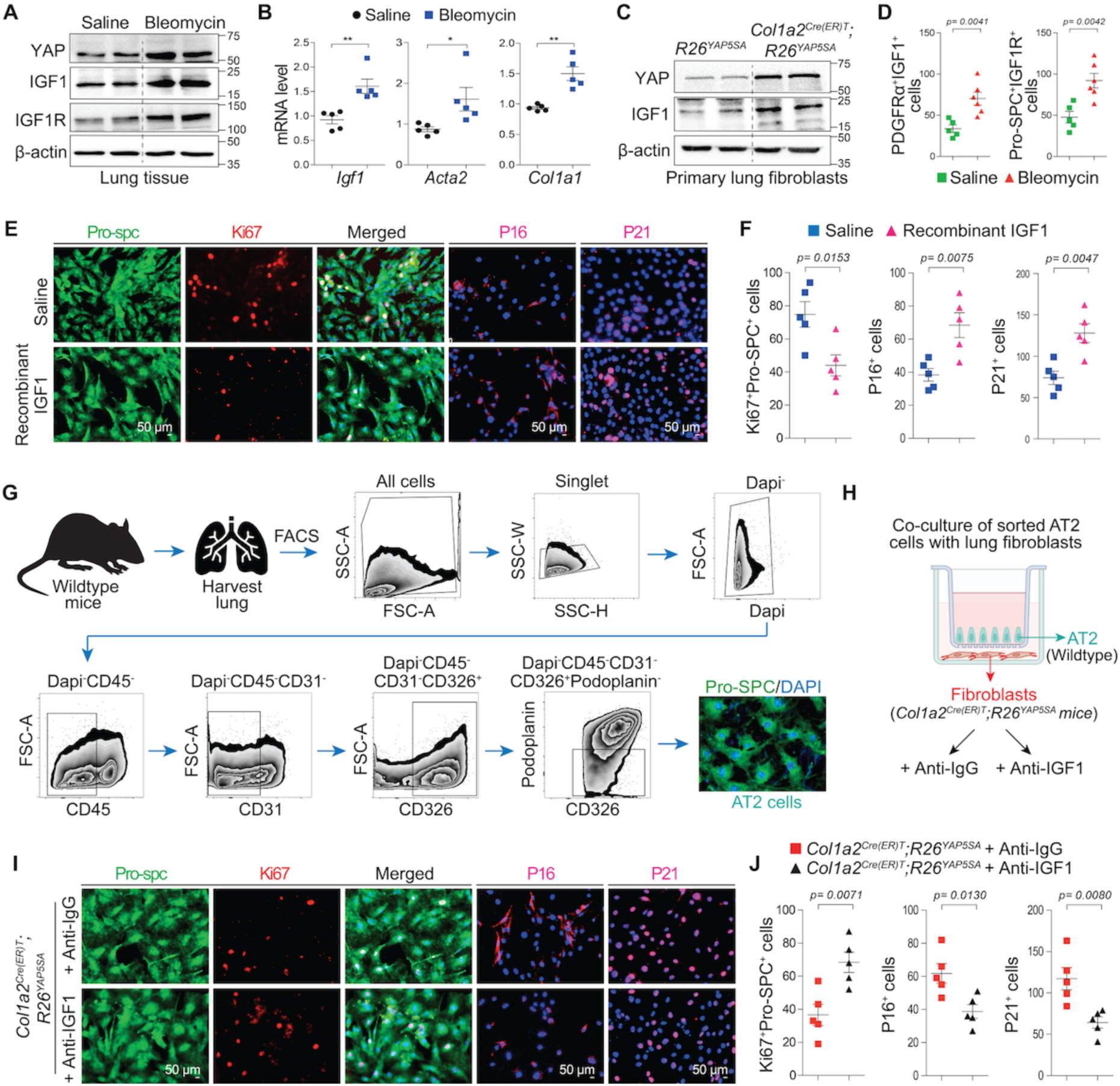
: YAP activity in fibroblasts affects AT2 cell homeostasis through IGF1-IGF1R-mediated paracrine signaling. (A) Immunoblot analysis of YAP, IGF1, and IGF1R on mouse lungs treated with saline or bleomycin for 14 days. Quantification of indicated proteins relative to β-actin. (B) Real-time qPCR for profibrotic genes *Acta2, Col1a1,* and *Igf1* on primary mouse lung fibroblasts isolated and cultured for 96 hours from 14 days saline or bleomycin treated mice (n = 5). (C) Immunoblot analysis of YAP and IGF1 on mouse primary lung fibroblasts from *R26^YAP5SA^*and *Col1a2^Cre(ER)T^; R26^YAP5SA^* mice challenged to intraperitoneal injection of tamoxifen (1 mg per kg body weight/day) for 5 alternative days before harvesting. (D) Quantification from immunofluorescence staining of IGF1 co-stained with PDGFRα, and immunofluorescence staining of IGF1R co-stained with Pro-SPC in wildtype mice lung sections at 14 days post-bleomycin injury compared to saline-treated controls (n = 5). (E-F) Immunostaining and quantification of Pro-SPC+Ki67+, or p16+ and p21+ AT2-cells (n=5) treated under IGF1(10ng/ml) or saline for 72 hours. (C-D) Graphical presentation for flow cytometry gating strategy to isolate AT2 cells from wild-type mice lungs defined by Dapi−CD45−CD31−CD326+Podoplanin− and subsequently co-cultured with lung fibroblasts from *R26^YAP5SA^* or *Col1a2^Cre(ER)T^; R26^YAP5SA^* lungs in the presence of IgG or IGF1-neutralizing antibody. (I-J) Immunostaining and quantification of Pro-SPC+Ki67+, or p16+ and p21+ AT2-cells from *R26^YAP5SA^* or *Col1a2^Cre(ER)T^; R26^YAP5SA^* lungs on co-culture insert where fibroblasts were exposed to IgG or IGF1-neutalizing antibody for 72 hours when cultured underneath of sorted AT2 cells (n=5). Data are presented as mean±SEM, with statistical comparisons made using a two-tailed unpaired t-test. Significance levels are indicated with p-value (significant at p<0.05).

### Verteporfin-mediated YAP/TAZ inhibition prevents fibrosis progression in bleomycin-challenged lungs

To test that inhibiting YAP/TAZ signaling could prevent or treat post-injury lung fibrosis, wild-type mice were treated with verteporfin (an FDA-approved YAP/TAZ inhibitor, 50 mg/kg) or DMSO every other day, beginning one day post-bleomycin administration and continuing until day 21 (Figure 7A). Lung fibrosis was assessed using Sirius Red staining, qRT-PCR, immunoblotting, and immunostaining. Verteporfin treatment markedly reduced fibrosis. Compared to controls, mice showed lower Ashcroft scores and hydroxyproline levels, along with decreased expression of profibrotic genes such as *Acta2, SM22α, Col1a1, P3h4, Plod2, Periostin, and Ccl2* (Figure 7B-E). Protein assays confirmed a reduction in Collagen I and α-SMA levels (Figure 7F-I). Notably, verteporfin preserved alveolar epithelial regeneration, with increased pro-SPC⁺ AT2 and Hopx⁺ AT1 cell populations in bleomycin-injured lungs (Figure 7J-K). To assess direct effects on fibroblasts, primary lung fibroblasts were treated with DMSO, TGF-β1 or TGF-β1 ± verteporfin (2 µM) for 96 h. TGF-β1 alone boosted expression of myofibroblast and collagen-processing genes (*Col1a1, Acta2, SM22α, Plod2, P3h4, Lox, Fkbp10*) and enhanced Collagen I and α-SMA stress fiber formation (Supplementary Fig 5A-B). However, co-treatment with verteporfin suppressed these profibrotic responses, halting fibroblast-to-myofibroblast transition and collagen production, demonstrating its direct anti-fibrotic effects on fibroblasts.

**Figure 7:**
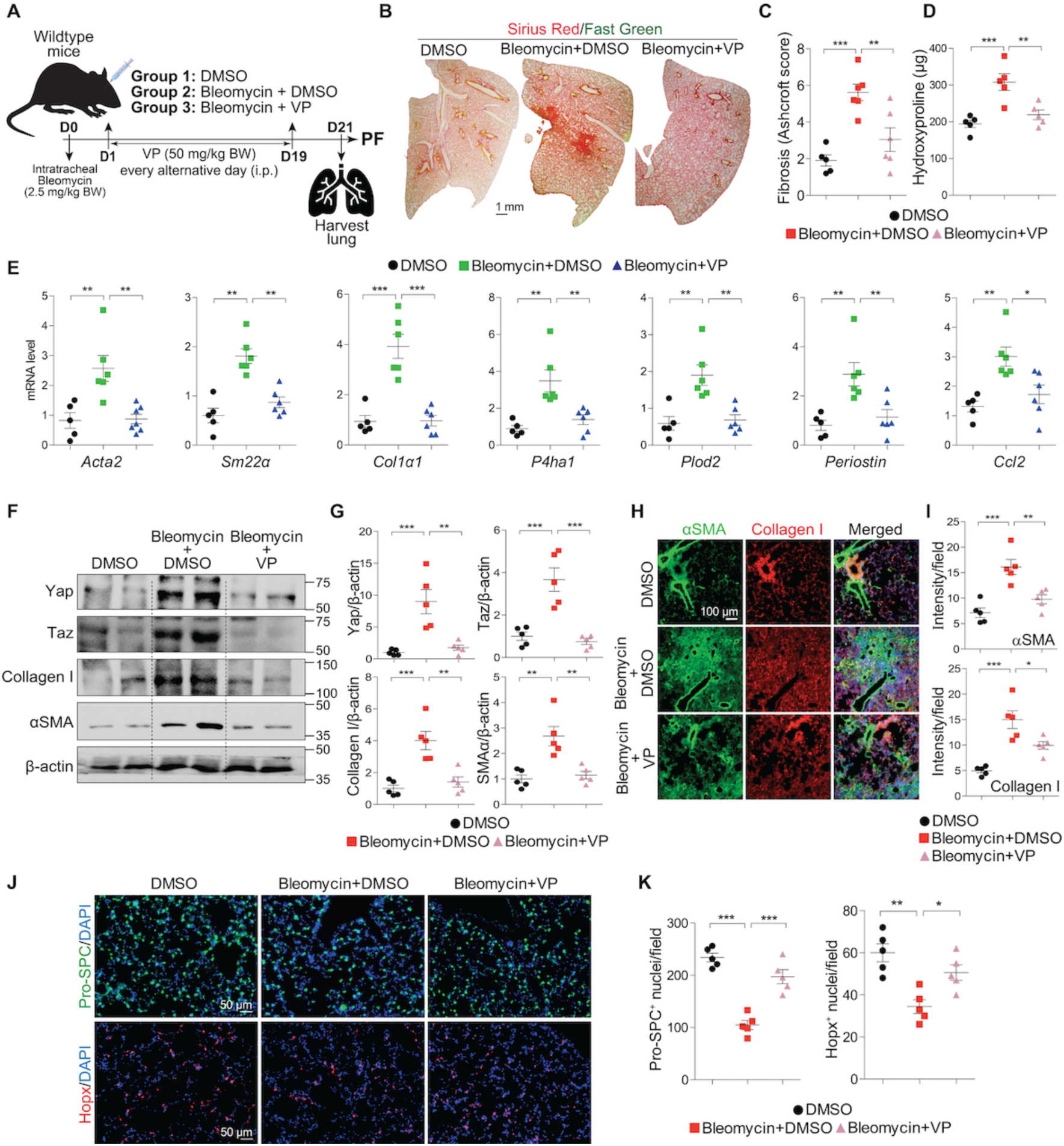
Verteporfin-mediated YAP/TAZ inhibition prevents fibrosis progression in bleomycin-challenged lungs. (A) Graphical presentation of experimental design; wild-type mice were challenged to one dose of intratracheal administration of bleomycin (2.5 mg/kg body weight) at day 0 followed by intraperitoneal injection of verteporfin (50 mg/kg body weight) or DMSO from day 1 until day 19, at every alternative day as indicated. DMSO was used as vehicle control. Lungs were harvested at 21 days post-bleomycin injury to assess the lung fibrosis. (B-C) Sirius Red/Fast Green staining and quantification (presented as Ashcroft score) on lung sections from wild-type mice (n = 5-6) treated with DMSO or bleomycin with/without verteporfin for 21 days. (D) Quantification of hydroxyproline content using lung tissue from wild-type mice (n = 5-6) treated with DMSO or bleomycin with/without verteporfin for 21 days. (E) Real-time qPCR for profibrotic genes *Acta2*, *SM22α*, *Col1a1, P4ha1, Plod2, Periostin* and *Ccl2* on lung tissue from wild-type mice (n = 5) treated with DMSO or bleomycin with/without verteporfin for 21 days. (F-G) Immunoblot analysis and quantification of Yap, Taz, collagen I and αSMA protein levels using lung tissue lysate from wild-type mice (n = 5) treated with DMSO or bleomycin with/without verteporfin for 21 days. Quantification of indicated proteins relative to β-actin. (H-I) Immunofluorescence staining and quantification of αSMA and collagen I in lung sections from wild-type mice (n = 5) treated with DMSO or bleomycin with/without verteporfin for 21 days. (J-K) Immunostaining and quantification of Pro-SPC+AT2-cells and Hopx+AT1-cells in wild-type mouse lungs (n=5) treated with DMSO or bleomycin with/without verteporfin for 21 days. Fluorescence intensity was quantified with imageJ software. The data are represented as the mean ± SEM; comparison by one-way ANOVA, followed by Tukey’s post-hoc test. *, *p* < 0.05; **, *p* < 0.01; ***, *p* < 0.001; NS, not significant.

### Verteporfin-mediated YAP/TAZ inhibition reverses established fibrotic lesions in bleomycin-challenged lungs

To assess verteporfin’s ability to reverse established fibrosis, we first induced lung fibrosis in wild-type mice using bleomycin and allowed fibrotic lesions to develop for 14 days before initiating therapy. Mice were then treated every other day with either vehicle (DMSO) or verteporfin (50 mg/kg body weight) until day 30 (Figure 8A). Verteporfin significantly attenuated existing fibrosis, reducing Ashcroft scores and hydroxyproline content while suppressing expression of fibrotic markers such as *Acta2*, *SM22α*, *Col1a1*, *P3h4*, *Plod2*, *Periostin*, *Ccl2* (Figure 8B-E). Importantly, immunoblot and histological analyses demonstrated verteporfin’s ability to degrade accumulated Collagen I and α-SMA deposits from injured lungs (Figure 8F-I). We also assessed whether verteporfin could protect against bleomycin-induced epithelial injury and enhance post-injury regeneration. Immunofluorescence for pro-SPC⁺ (AT2) and Hopx⁺ (AT1) cells in verteporfin-treated lungs showed increased epithelial cell counts after injury (Figure 8J-K). To assess verteporfin’s direct antifibrotic effects, we stimulated lung fibroblasts with TGFβ1 for 96 hours before initiating verteporfin treatment. While TGFβ1 robustly upregulated myofibroblast markers (*Col1a1, Acta2, SM22a*) and collagen-modifying enzymes (*Plod2, P3h4, Lox, Fkbp10*), subsequent verteporfin treatment effectively reversed this fibrogenic activation (Supplementary Figure 6A). Immunofluorescence analysis confirmed that verteporfin significantly reduced TGFβ1-induced Collagen I production and α-SMA stress fiber formation (Supplementary Figure 6B), demonstrating its capacity to counteract established profibrotic signaling in lung fibroblasts.

**Figure 8:**
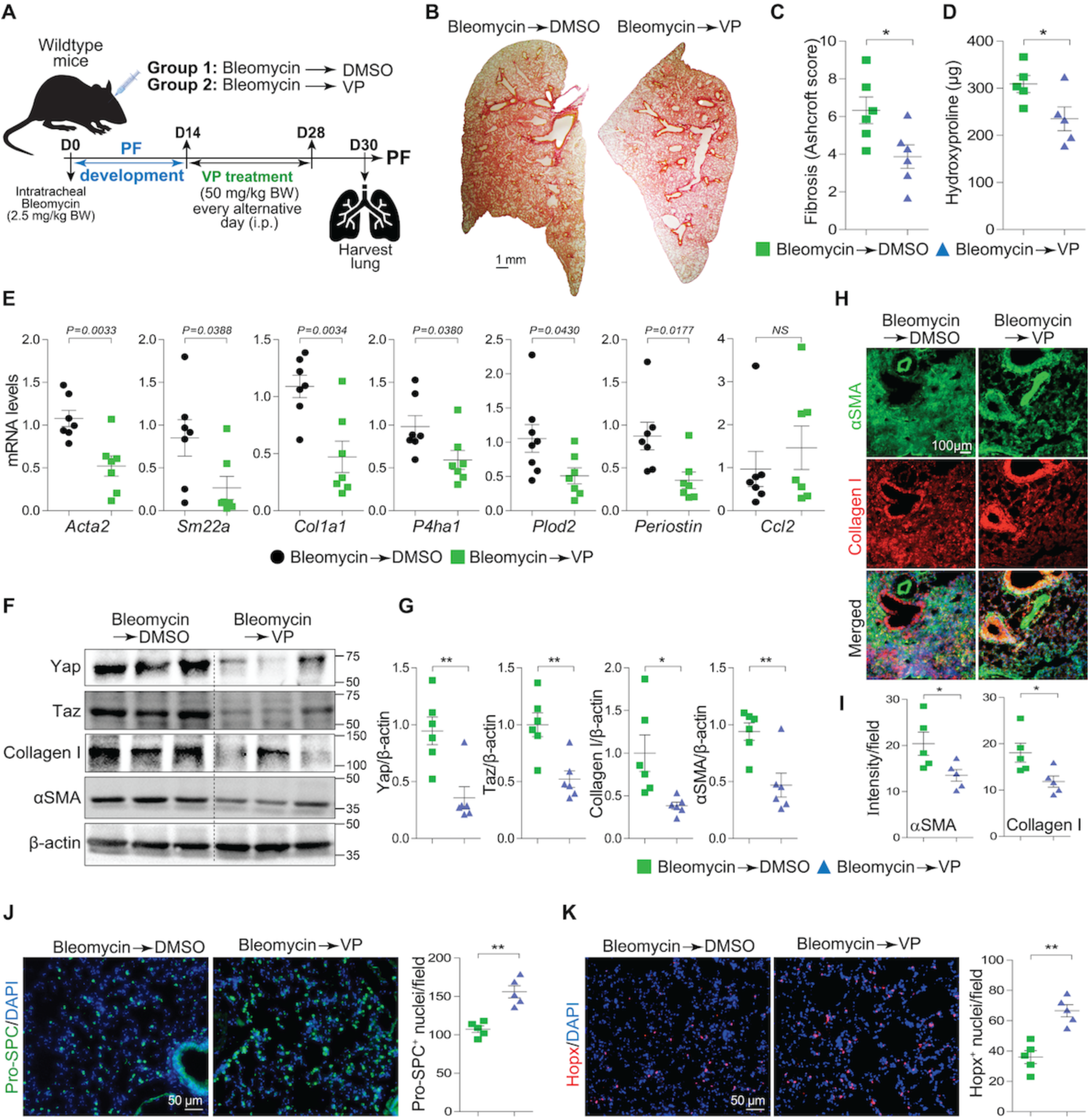
Verteporfin-mediated YAP/TAZ inhibition reverses established fibrotic lesions in bleomycin-challenged lungs. (A) Graphical presentation of experimental design; wild-type mice were challenged to one dose of intratracheal administration of bleomycin (2.5 mg/kg body weight) at day 0 followed by intraperitoneal injection of verteporfin (50 mg/kg body weight) or DMSO started from day 14 until day 28. Administration of DMSO/verteporfin was performed at every alternative day as indicated. Lungs were harvested at 30 days post-bleomycin injury to assess the lung fibrosis. (B-C) Sirius Red/Fast Green staining and quantification (presented as Ashcroft score) on lung sections from wild-type mice (n = 6) challenged to bleomycin for 14 days and then post-treated with verteporfin or DMSO for 14 days. (D) Quantification of hydroxyproline content using lung tissue from wild-type mice (n = 5) challenged to bleomycin for 14 days and then post-treated with verteporfin or DMSO for 14 days. (E) Real-time qPCR for profibrotic genes *Acta2*, *SM22α*, *Col1a1, P4ha1, Plod2, Periostin* and *Ccl2* on lung tissue from wild-type mice (n = 7) challenged to bleomycin for 14 days and then post-treated with verteporfin or DMSO for 14 days. (F-G) Immunoblot analysis and quantification of Yap, Taz, collagen I and αSMA protein levels using lung tissue lysate from wild-type mice (n = 6) challenged to bleomycin for 14 days and then post-treated with verteporfin or DMSO for 14 days. Quantification of indicated proteins relative to β-actin. (H-I) Immunofluorescence staining and quantification of αSMA and collagen I in lung sections from wild-type mice (n = 5) challenged to bleomycin for 14 days and then post-treated with verteporfin or DMSO for 14 days. (J-K) Immunostaining and quantification of Pro-SPC+AT2-cells and Hopx+AT1-cells in lung sections from wild-type mice (n = 5) challenged to bleomycin for 14 days and then post-treated with verteporfin or DMSO for 14 days. Fluorescence intensity was quantified with imageJ software. The data are represented as the mean ± SEM; comparison by one-way ANOVA, followed by Tukey’s post-hoc test. *, *p* < 0.05; **, *p* < 0.01; ***, *p* < 0.001; NS, not significant.

### Verteporfin enhances Collagen I degradation by modulating MMP/TIMP balance and suppressing YAP/TAZ-driven pro-inflammatory signaling

Given the established roles of matrix metalloproteinases (MMPs) and their tissue inhibitors (TIMPs) in Collagen I degradation, we hypothesized that verteporfin may modulate MMP and TIMP expression to enhance Collagen I turnover by myofibroblasts. To investigate how verteporfin regulates Collagen I turnover in a pro-fibrotic environment, primary lung fibroblasts were pre-treated with TGF-β1 for 96 hours, followed by exposure to verteporfin (2 μM) for the indicated time points and subsequent analysis by immunoblotting. Verteporfin treatment resulted in a marked reduction in YAP, TAZ, and Collagen I protein levels following TGF-β1 stimulation (Figures 9A-B). To assess how verteporfin influences collagen degradation, we analyzed the pro- and active forms of MMP1, MMP3, and MMP9. TGF-β1 stimulation increased pro-MMP1 (∼52 kDa) without affecting its active form (∼22 kDa). Verteporfin treatment, however, reduced pro-MMP1 while maintaining active MMP1 levels, leading to a sustained increase in the active/pro-MMP1 ratio from 15 minutes to 6 hours. A similar pattern was observed for MMP9 where TGF-β1 upregulated pro-MMP9 (∼92 kDa) but not active MMP9 (∼40 kDa), whereas verteporfin decreased pro-MMP9 while enhancing both the ∼40 kDa and ∼27 kDa active isoforms, significantly raising the active/pro-MMP9 ratio between 1 and 6 hours (Figures 9C-F). In contrast, verteporfin suppressed both pro- and active MMP3 in a time-dependent manner, suggesting divergent regulation among these MMPs (Supplementary Figure 7). Verteporfin further attenuated TGF-β1-induced TIMP1 and TIMP2 expression in a time-dependent manner (Figures 9C-F). This reduction in TIMP levels, coupled with enhanced MMP1 and MMP9 activation, likely promotes Collagen I degradation in a TGFβ1-rich fibrotic environment.

**Figure 9:**
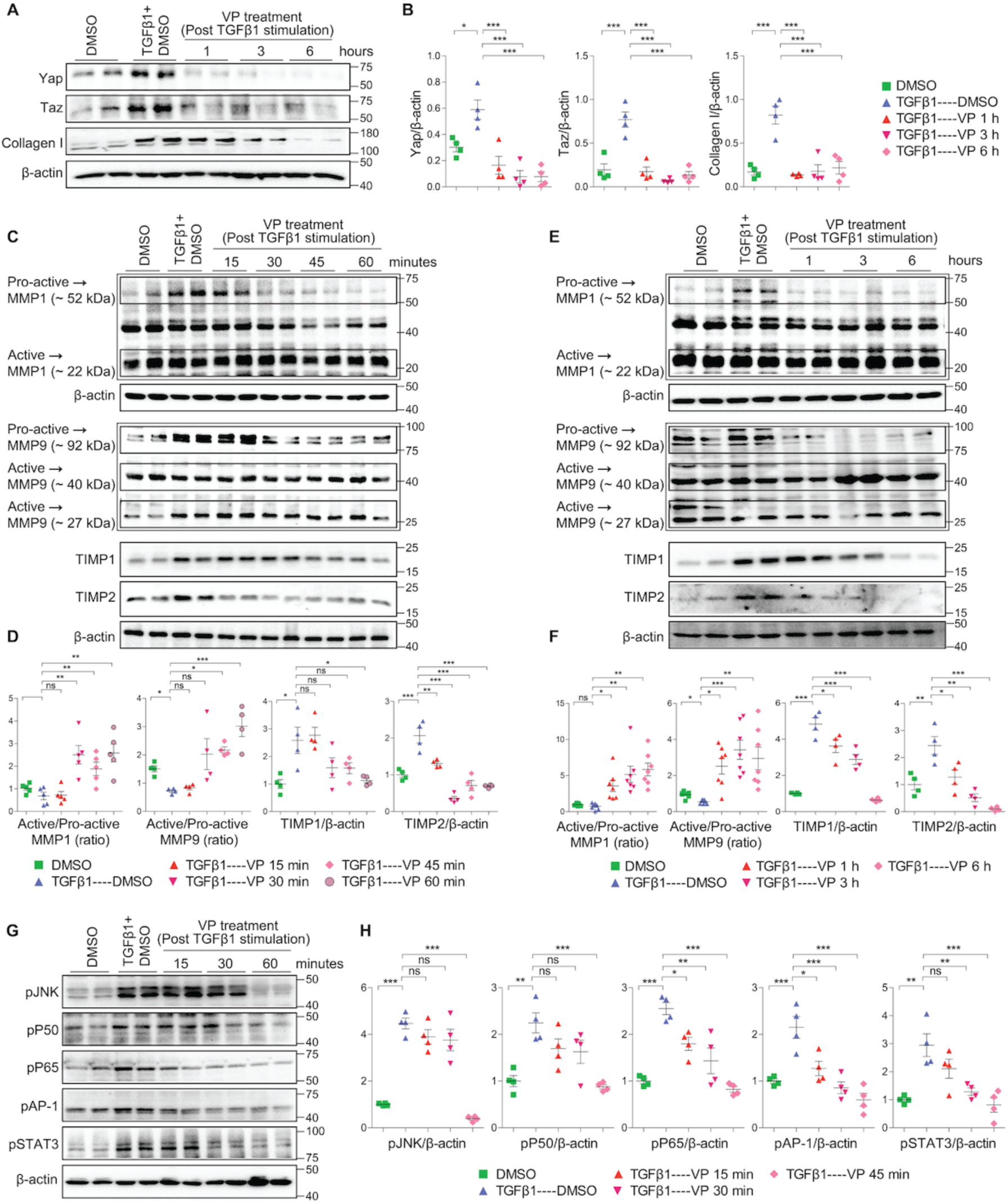
Verteporfin affects Collagen I remodeling by modulating MMP/TIMP balance and suppressing YAP/TAZ driven pro-inflammatory signaling. Primary mouse lung fibroblasts were cultured for 96 hours in the presence of TGFβ1, followed by a post-treatment with VP for 15, 30, 45 and 60 minutes respectively. In a separate experiment, post-treatment with VP in the presence of TGFβ1 was also performed for 1, 3 and 6 hours respectively. DMSO was used as vehicle control. (A-B) Immunoblot analysis and quantification of Yap, Taz, and collagen I protein levels using lung fibroblasts treated with TGFβ1 for 96 hours followed by post-treatment with VP as indicated time points. (C-F) Immunoblot analysis and quantification of MMP1 and MMP9 protein levels marked as pro-active and active forms together with TIMP1 and TIMP2 proteins using lung fibroblasts treated with TGFβ1 for 96 hours followed by post-treatment with VP for as indicated time points. (G-H) Immunoblot analysis and quantification of pJNK, pP50, pP65, pAP1, and pSTAT3 protein levels using lung fibroblasts treated with TGFβ1 for 96 hours followed by post-treatment with VP as indicated time points. Quantification of indicated proteins relative to β-actin. The data are represented as the mean ± SEM; comparison by one-way ANOVA, followed by Tukey’s post-hoc test. *, *p* < 0.05; **, *p* < 0.01; ***, *p* < 0.001; NS, not significant.

**Figure 10:**
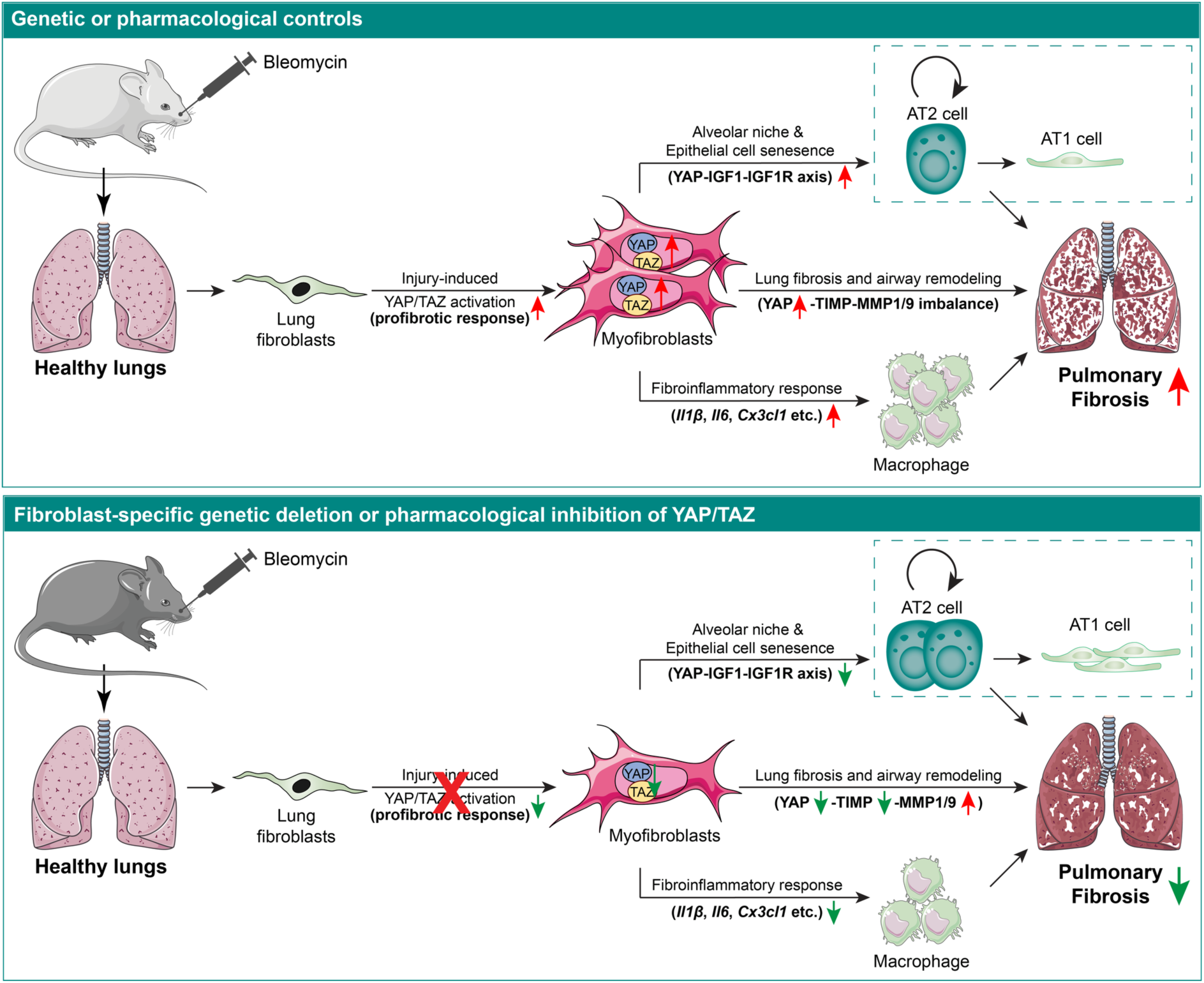
Graphical model demonstrating fibroblast-specific role of YAP/TAZ in pulmonary fibrosis. The schematic illustrates YAP/TAZ activation in fibroblasts during bleomycin-induced lung fibrosis, driving fibrotic progression through multiple mechanisms. YAP/TAZ promote ECM remodeling via the TIMP-MMP1/9 axis and enhance fibroinflammatory responses by regulating cytokine/chemokine production and macrophage recruitment. Additionally, fibroblast-derived YAP activation disrupts alveolar epithelial homeostasis by inducing senescence through IGF1-IGF1R paracrine signaling. Genetic ablation of *Yap/Taz* in fibroblasts or pharmacological inhibition (e.g., verteporfin) attenuates myofibroblast activation, reduces fibrosis, reduces inflammation and macrophage homing, and promotes alveolar regeneration post-injury.

To elucidate the molecular mechanisms through which YAP/TAZ influences fibro-inflammatory responses, we assessed the activation status of key pro-inflammatory transcription factors in primary lung fibroblasts. Western blot analysis revealed robust activation of phosphorylated JNK (pJNK), NF-κB subunits (pP50, pP65), phosphorylated AP-1 (pAP-1), and phosphorylated STAT3 (pSTAT3) following 96 hours of TGF-β1 stimulation. Verteporfin treatment inhibited the activation of these factors in a time-dependent manner (Figures 9G-H), suggesting that YAP/TAZ inhibition disrupts pro-inflammatory pathways central to pulmonary fibrosis progression and exacerbation.

## Discussion

Idiopathic pulmonary fibrosis is a progressive and fatal lung disease characterized by dysregulated extracellular matrix (ECM) remodeling, leading to irreversible scarring and respiratory failure. Central to IPF pathogenesis are myofibroblasts, differentiated fibroblasts that drive fibrosis through excessive ECM production and tissue contraction. Myofibroblasts further exacerbate disease progression by secreting pro-fibrotic mediators that recruit and polarize macrophages into a profibrotic phenotype, while also inducing alveolar epithelial cell (AEC) apoptosis and senescence. This triad of interactions myofibroblast-macrophage crosstalk and myofibroblast-mediated epithelial dysfunction creates a vicious cycle of persistent injury, impaired repair, and pathological remodeling. Therefore, identifying the molecular mechanisms governing fibroblast activation and deactivation is critical for developing targeted therapies for pulmonary fibrosis (PF/IPF). In this study, we identified hyperactivation of YAP/TAZ in fibroblasts from PF patient lungs and bleomycin-injured mouse lungs. Genetic deletion or pharmacological (verteporfin) inhibition of fibroblast-specific YAP/TAZ signaling not only prevented pathological fibroblast-to-myofibroblast transition but also reversed established fibrosis.

Recent studies have demonstrated the contribution of Hippo signaling effectors YAP and TAZ in in pulmonary fibrosis pathogenesis (11–17). While these transcriptional regulators are markedly upregulated in fibrotic lungs of IPF patients, their functional impact exhibits striking cell-type specificity, driving distinct profibrotic mechanisms across different cellular compartments (12, 14). For example, While YAP/TAZ in IPF-derived and TGF-β1-activated fibroblasts critically drive myofibroblast differentiation and pathological ECM deposition *in vitro*, their systemic siRNA-mediated knockdown aggravated bleomycin-induced pulmonary fibrosis (13). This apparent contradiction likely stems from the distinct, context-dependent roles of YAP/TAZ in alveolar epithelial cells versus fibroblasts as in contrast to fibroblasts, YAP/TAZ activation in epithelial cells has be shown to be critical for resolving lung inflammation and driving epithelial regeneration after injury (11–13, 16). Combined knockdown of YAP and TAZ significantly reduces the expression of contractile genes such as *Acta2*, *Cnn1*, and *Tagln* in lung fibroblasts from IPF patient and TGF-β1-activated fibroblasts (11). Notably, dual knockdown of YAP/TAZ more effectively suppresses fibroblast contractility than individual knockdowns, reflecting the redundant yet cooperative roles of these transcriptional cofactors in promoting fibrogenic function (12, 14). While *in vitro* studies demonstrate that combined YAP/TAZ knockdown effectively suppresses fibroblast activation, the *in vivo* consequences of fibroblast-specific dual deletion remain unexplored, particularly in the context of pulmonary fibrosis. To address this, we generated fibroblast-specific YAP/TAZ double-knockout mice (*Col1a2^Cre(ER)T^;Yap^flox/flox^;Taz^flox/flox^*) to systematically investigate how their coordinated loss impacts fibroinflammatory responses, myofibroblast differentiation, and alveolar regeneration/repair post bleomycin-induced injury. Fibroblast-specific YAP/TAZ deletion significantly attenuated bleomycin-induced pulmonary fibrosis, shown by reduced hydroxyproline content and lower collagen I deposition in lung tissue. These results align with previous findings from whole-body *Taz*-heterozygous mice, which also exhibited resistance to bleomycin-induced PF (18). Fibroblast-specific overexpression of YAP (using *Col1a2^Cre(ER)T^;R26^YAP5SA^* mice) exacerbated bleomycin-induced fibroinflammatory and fibrogenic responses, reinforcing previous observations in IPF fibroblasts that YAP activation alone is sufficient to drive pulmonary fibrosis. Complementing genetic evidence, pharmacological modulation of YAP/TAZ signaling demonstrated consistent anti-fibrotic efficacy across experimental systems. Pharmacological inhibition of YAP/TAZ signaling in DRD1-positive lung fibroblasts using the selective DRD1 agonist dihydrexidine successfully attenuated fibrotic progression (13). This intervention not only reversed established bleomycin-induced fibrosis *in vivo* but also suppressed pathological activation of IPF patient-derived lung fibroblasts *in vitro*, demonstrating cross-species therapeutic potential. A recent study found that statins can restrict YAP nuclear localization and attenuate established lung fibrosis in bleomycin models (19). Our present study demonstrated that verteporfin-mediated inhibition of YAP/TAZ not only halted the development and progression of fibrosis but also reversed established fibrotic lesions, highlighting its therapeutic potential across all stages of the disease. This convergence of findings, from genetic YAP/TAZ knockout and overexpression mouse models to pharmacological drugs like verteporfin, statins, and dihydrexidine strongly validates YAP/TAZ as central regulators of fibrosis that are amenable to clinical targeting across diseases. Therapeutic synergy between FDA-approved drugs and novel targeted agents may offer superior clinical outcomes for IPF patients. By combining existing anti-fibrotics (e.g., nintedanib/pirfenidone) with emerging YAP/TAZ inhibitors like verteporfin, we could simultaneously target multiple pathogenic pathways addressing both matrix deposition and mechanotransduction-driven fibrogenesis. This combinatorial approach may enhance efficacy while potentially reducing individual drug dosages and associated side effects.

Fibroblasts play a pivotal role in orchestrating the fibroinflammatory response following lung injury through their secretion of cytokines/chemokines. These mediators regulate both the recruitment and phenotypic polarization of inflammatory cells while influencing their survival within injured tissue. The marked downregulation of IL-1β, IL-6, Cx3cl1, Cx3cr1 and TNF-α observed in fibroblast-specific YAP/TAZ knockout lungs following bleomycin injury is particularly significant given their established roles in pulmonary fibrosis. For example, IL-1β drives acute pulmonary inflammation by recruiting neutrophils and macrophages (20, 21). Its chronic expression induces sustained TGF-β1 and PDGF production, promoting fibrosis through collagen deposition and myofibroblast expansion (22). IL-6 promotes fibroblast proliferation and collagen deposition via TGF-β pathway activation and elevated IL-6 correlates with lung function decline in PF patients (23, 24). Similarly, Cx3cl1-Cx3cr1 chemokine-receptor pair regulates myeloid cell dynamics and modulate fibroblast activation (25, 26). Our findings align with a growing body of evidence that connects YAP/TAZ to the regulation of cytokine expression and immune responses across various pathological conditions (10, 27–36). Our study showed a blunted fibroinflammatory response characterized by reduced F4/80+ macrophage infiltration and diminished secretion of inflammatory cytokines/chemokines, demonstrating their causal role in PF pathogenesis.

IPF is marked by accumulation of fibrillar collagens, with type I collagen being the most abundant extracellular matrix protein. Pathogenic myofibroblasts drive this accumulation through excessive synthesis and insufficient degradation of collagen I, an imbalance that is a hallmark of PF/IPF (37, 38). In fibrosis, restoring physiological collagen I degradation requires coordinated MMPs activation and TIMP modulation. MMP-1 initiates the process by cleaving intact fibrillar collagen, while MMP-9 degrades the resulting fragments. Since TIMPs naturally regulate this proteolytic cascade, therapeutic strategies that rebalance MMP-TIMP activity could selectively reduce pathological collagen deposition in IPF (39–41). Our present study demonstrated that TGF-β1-induced profibrotic conditions activate YAP/TAZ in fibroblasts. Pharmacological inhibition of YAP/TAZ with verteporfin significantly promoted collagen I clearance through a dual regulatory mechanism, suppressing TIMP1/2 inhibition while enhancing MMP1/9 activation, evidenced by increased active/pro-active form ratios). These observations align with established antifibrotic mechanisms of current FDA approved IPF drugs Nintedanib and Pirfenidone. Nintedanib enhances pro-MMP-2 activity while reducing TIMP-2 levels in IPF fibroblasts, resulting in decreased TGF-β-driven collagen secretion and slowed disease progression (42). Pirfenidone suppresses TGF-β1–induced TIMP-1 expression and promotes MMP-2/9 activity, restoring ECM degradation pathways (43). Nintedanib mediates its anti-fibrotic effects on lung fibroblast, in part, by inhibiting TBK1 phosphorylation, thereby preventing nuclear translocation of the YAP/TAZ transcriptional complex (44). Selective inhibition of fibroblast YAP/TAZ signaling via the Dopamine Receptor D1 agonist dihydrexidine (DHX) attenuated fibrotic progression by enhancing extracellular matrix degradation. DHX stimulates cathepsin K-dependent collagen I breakdown and enhances MMP14 expression, switching fibroblasts from collagen-producing to matrix-resorbing cells and reducing tissue stiffness in lung fibrosis models (13). The anti-fibrotic potential of enhanced matrix degradation is well-established across multiple models. For example, macrophage-derived MMP9 overexpression attenuates bleomycin-induced PF, while IL-1β exerts protective effects in human lung fibroblasts by elevating MMP1/9-mediated collagenolysis (21, 45, 46). Similarly, proteasome inhibition reduces fibrosis in dermal fibroblasts through coordinated suppression of collagen I/TIMP1 and activation of MMP1, suggesting this approach may be therapeutic for other fibrotic conditions such as systemic sclerosis (47, 48). Our studies further revealed that verteporfin disrupts TGF-β1-driven expression of key collagen-modifying enzymes (*Plod2, P3h4, Fkbp10*) at the transcriptional level. These enzymes critically regulate collagen crosslinking and maturation, and their pathological overexpression contributes to aberrant ECM architecture and tissue stiffness in PF (49, 50). Notably, Fkbp10 knockdown alone reduces collagen secretion, supporting that verteporfin’s suppression of these enzymes impairs post-translational collagen processing (50). Combined with its MMP-activating effects, these data demonstrate verteporfin’s dual capacity to modulate collagen I homeostasis, simultaneously inhibiting pathological synthesis while promoting degradation-thereby addressing both arms of ECM dysregulation in pulmonary fibrosis.

Fibroblast-mediated fibroinflammatory responses critically impair lung regeneration by disrupting alveolar epithelial cell differentiation and function (51–54). In pulmonary fibrosis, the loss or dysfunction of AT1 and AT2 cells following injury drives disease progression and respiratory decline (54–56). As the primary progenitor population, AT2 cells normally proliferate and differentiate into AT1 cells to restore alveolar integrity post-injury, a process supported by specialized alveolar niche fibroblasts that promote AT2 self-renewal and differentiation (52, 57–59). However, pathological transformation of fibroblasts into hyperactivated myofibroblasts disrupts this regenerative niche, suppressing AT2 cell function and creating a self-perpetuating cycle of fibrotic progression (57, 58, 60, 61). Our genetic experimental models demonstrated that this balance is critically regulated by YAP/TAZ signaling, as genetic deletion or pharmacological inhibition (verteporfin) enhanced alveolar epithelial differentiation and function, whereas fibroblast-specific YAP overexpression impaired repair. This opposing epithelial cell phenotype likely stems from differential effects on myofibroblast activation where YAP/TAZ deletion/inhibition preserves the regenerative niche by suppressing pathological myofibroblast transformation, and YAP hyperactivation disrupts it through excessive myofibroblast accumulation. To determine whether the epithelial effects were mediated solely by niche improvement or involved direct signaling, we employed co-culture systems. These experiments revealed that YAP-activated myofibroblasts directly suppress epithelial cell proliferation and promote senescence. Mechanistically, we identified that these myofibroblasts secrete elevated Insulin-like growth factor 1 (IGF1), which activates IGF1 receptors on alveolar epithelial cells to drive these dysfunctional states, including senescence. Neutralization of IGF1 with blocking antibodies restored epithelial proliferation and reduced senescence markers, confirming the functional significance of this pathway. IGF1 is emerging as a key mediator of fibrogenesis and cellular senescence across tissues including lung where IGF1 levels are elevated in lungs of IPF patients and bleomycin-treated mice (62–68). IGF1 exerts time-dependent effects on AT2 cells and fibrosis progression. In the short term, IGF1 supports lung repair by promoting AT2 cell proliferation and expansion during acute injury, contributing to alveolar regeneration (69, 70). However, prolonged IGF1 signaling leads to chronic activation of IGF1R and increases of senescence markers, driving AT2 cells into irreversible growth arrest and impairing their differentiation into AT1 cells, ultimately disrupting alveolar repair and maintaining fibrotic remodeling (65, 68, 71, 72). Consistently, alveolar epithelial cells-specific IGF1R knockout mice (*SftpcCre/ERT2;Igf1r^flox/flox^*) markedly reduced AT2 cell senescence (66). This paracrine YAP-IGF1-IGF1R axis represents a direct cell-to-cell communication pathway through which profibrotic myofibroblasts actively impair epithelial regeneration. These results establish that YAP/TAZ promote lung repair through two interdependent mechanisms, creating a permissive microenvironment while simultaneously facilitating direct fibroblast-epithelial cell communication.

In summary, our study demonstrated that fibroblast YAP/TAZ activation plays a pivotal role in pulmonary fibrosis pathogenesis by orchestrating three key pathological processes: promoting fibroblast-to-myofibroblast differentiation and excessive ECM deposition, modulating fibroinflammatory response, and disrupting alveolar regeneration. Through both genetic and pharmacological approaches, we showed that selective inhibition of fibroblast YAP/TAZ signaling not only attenuates these profibrotic mechanisms but also actively promotes lung repair. These findings suggests that targeting this pathway could achieve comprehensive therapeutic effects unmatched by current single-mechanism approaches in IPF treatment.

## Materials and methods

### Mice

Fibroblast-specific *Yap/Taz*-double knockout mice were generated by the *Col1a2^Cre(ER)T^* transgenic mice crossing with *Yap^flox/flox^; Taz^flox/flox^* mice (73, 74). The resulting *Col1a2^Cre(ER)T^; Yap^flox/+^; Taz^flox/+^* offspring were then backcrossed to *Yap^flox/flox^_;_ Taz^flox/flox^* mice to generate *Col1a2^Cre(ER)T^; Yap^flox/flox^; Taz^flox/flox^* mice. In *Yap/Taz* genetic deletion experiments, *Yap^flox/flox^; Taz^flox/flox^,* mice were used as controls and *Col1a2^Cre(ER)T^; Yap^flox/flox^; Taz^flox/flox^* as *Yap/Taz* double knockout mice (*dKO*). *Yap* and *Taz* floxed mice were genotyped as described previously (73, 74). Likewise, for *Yap* overexpression in lung fibroblast, *Col1a2^Cre(ER)T^* mice were crossed with *Rosa26^Yap5SA/Yap5SA^* knock-in strain to obtain *Col1a2^Cre(ER)T^; Rosa26^Yap5SA^* mice. *Rosa26^Yap5SA^*mice we used as control for overexpression study. *Rosa26^Yap5SA^*mice were genotyped as described previously (75). For *Yap/Taz* deletion in fibroblasts, both control and *dKO* mice were subjected to intraperitoneal injection of 1 mg per kg body weight of tamoxifen (Sigma-Aldrich, catalogue no. T5648) for five alternative days from the first day of bleomycin instillation to induce Cre-mediated recombination. Similarly, for *Yap* activation in lung fibroblast, both *R26^Yap5SA^* and *Col1a2^Cre(ER)T^; R26^Yap5SA^* mice were injected with tamoxifen as described above. Littermate controls were used in all experiments. All animal procedures were approved by the Institutional Animal Care and Use Committee at Duke-NUS Medical School/Singhealth conforming to the Guide for Care and Use of Laboratory Animals (National Academies Press, 2011). All mice were maintained on a mixed (C57BL/6 and Sv/129) genetic background. Both male and female mice aged from 12-16 weeks were used for analysis.

### Human samples

Adult human control lung tissues were purchased from Novus Biologicals (Catalogue no. NBP2-77573, NBP2-30182 and NBP2-42696) and human lung tissues diagnosed with fibrotic diseases were purchased from Origene (Catalogue no. CS504154, CS600794 and CS505314). Frozen lung tissue sections containing slides were stored in −80°C before use and performed immunostaining assay as required.

### Bleomycin induction of pulmonary fibrosis

*Yap^flox/flox^;Taz^flox/flox^*, *Col1a2^Cre(ER)T^; Yap^flox/flox^; Taz^flox/flox^*, *Rosa26^Yap5SA^* and *Col1a2^Cre(ER)T^; R26^Yap5SA^* mice were asleep with Isoflurane (Piramal, USA) followed by intratracheal administration of bleomycin (Abcam, catalog no. ab142977) as 2.5 mg/kg body weight. To administer the bleomycin, the tongue of mice was retracted and intubated a 22-gauge IV catheter (B. Braun, catalog no. 4251628-03) into the trachea and quickly instilled the bleomycin. Same amount of saline was administered to group the controls animals against bleomycin injury. The mice were then euthanized on day 7 or 14 to evaluate the bleomycin-induced pulmonary fibrosis. Broncho alveolar lavage (BAL) fluid was obtained by administration of 2.5 ml of saline supplemented with 100μM EDTA and protease-phosphatase inhibitor cocktail (Sigma, catalog no. 11836170001 and 4906845001) and subsequently perfused lungs were harvested and processed for further analysis including gene expression, histological or immunoblot analysis. In other experiments of fibrosis analysis, verteporfin (sigma) (50 mg/kg BW) was administered from the next day of bleomycin instillation in regard to preventive study or 14 days post bleomycin treatment in regards to reversal study. DSMO was administered into the mice as control group of verteporfin. The left lung and the superior lobe of right lung were used for histological analysis; the middle lobe of right lung used for qRT-PCR analysis; the inferior lobe of right lung used for hydroxyproline assay and the post-caval lobe was used for western blot analysis. However, for flow cytometry analysis whole lung was used to sort and culture the alveolar epithelial type-II cells.

### Cell culture

#### Primary Lung Fibroblasts

Primary lung fibroblasts were isolated from mice as described earlier (76). Fibroblasts were cultured at 37°C and 5% CO2 in DMEM supplemented with 10% FBS, 1% penicillin/streptomycin. Following treatment conditions were used as time dependent manner to manipulate lung fibroblasts activation: recombinant mouse TGFβ1 (10 ng/mL, Biolegend, catalogue no. 763102) and verteporfin (2 μM, Sigma), TGFβ1 were resuspended in water whereas verteporfin was solubilized in DMSO (Sigma). To perform the qRT-PCR and western blot analysis, lung fibroblasts were seeded in 6-well cell culture plate. To conduct the immunostaining on cells, cultured fibroblasts were seeded in 24-well cell culture plate.

#### Primary Lung AT2 cells

Primary lung AT2 cells were isolated from mice using flow cytometry identified as Dapi−CD45−CD31−CD326+Podoplanin− cells. AT2 cells were grown for 14 days at 37°C and 5% CO2 in alveolar epithelial cell medium purchased from Sciencell (Catalog No. 3201). Cultured AT2 cells were then seeded into 12 mm Transwell with 0.4 µm pore polyester membrane inserts (Corning, Catalog No. 3460) to perform co-culture experiment lung fibroblasts. In co-culture experiment, fibroblasts were seeded on the underside of a Transwell ® insert. In the recombinant IGF1 (R&D, Catalog No. 791-MG-050) or neutralization of IGF1 (R&D, Catalog No. MAB791-100) experiments 10 ng/ml or 1 μg/ml were added to the medium for 72 hours respectively.

### Histology and immunohistochemistry

Histology and immunohistochemistry procedures were followed as described before (77). Briefly, lung lobes were fixed in 4% paraformaldehyde overnight at 4°C and subsequently performed paraffin embedding. Paraffin embedded 5 µm lung sections were subjected to perform Masson’s trichrome, Sirius Red staining, and other immunostaining. To evaluate the lung fibrosis the mean Ashcroft score was calculated as described previously (78). For immunostaining, lung sections were permeabilized with 0.1% Triton-X-100 in PBS and consequently performed antigen retrieval with antigen unmasking solution (Vector Laboratories, USA). Endogenous peroxidase activity was blocked with 3% hydrogen peroxide for 10 minutes and then sections were blocked for 1.5 hours using 5% bovine serum albumin (BSA) (Boval BioSolutions, Catalog no. LY-0080) at RT. Next, sections were incubated with specific primary antibodies diluted in 5% BSA solution at 4°C overnight followed by washing and subsequently probed with secondary antibodies from Invitrogen (Alexa Fluor 488, Alexa Fluor 568, Alexa Fluor 647 conjugated) diluted in 5% BSA blocking buffer for 2 hours at RT. Sections were washed and incubated with Dapi (Sigma, Catalog no. D9542) for 15 minutes. After washing sections were mounted with 100% glycerol and a staining pattern was visualized with a Leica fluorescence microscope.

To perform the Sirius Red staining lung sections were deparaffinized and hydrated with distilled water. Sections were then incubated with Picro-Sirius Red solution (Abcam, catalog no. ab246832) for one hour and subsequently washed with 1% Acetic acid followed by dehydration using ethanol and mounted with Depex (Sigma) for imaging. Likewise to perform Masson’s trichrome staining, sections were deparaffinized and fixed with Bouin’s solution (Sigma, catalog no.HT10132) for 30 minutes in a water bath at 58°C. Then, a mixture of Wiegert’s Solution A and B (Merck Millipore, catalog no.1.15973.0002) was applied in a ratio of 1:1 for 15 minutes followed by thoroughly washing the sections with distilled water. Next, 1% Ponceau Biebrich Scarlet solution (Sigma catalog no. B6008) and 1% Fuchsin acid solution (Merck Millipore 1.05231.0025) were applied for 5 minutes followed by washing the sections with distilled water. Then, the mixture of 2% Phosphomolybdic acid solution (Merck Millipore, catalog no. 1.00532.0025) and 2% Phosphotungstic acid solution (Merck Millipore, catalog no. 1.00583.0100) was added for 10 minutes at RT. Lastly, 2% methyl blue (Sigma, catalog no. M5528) was applied to the sections and incubated for 15 minutes followed by washing with 1% acetic acid.

To perform immunofluorescence on cells, we followed fixation with methanol/acetone for 5 min at −20°C. Next, cells were washed and incubated with indicated antibodies diluted in PBS containing 2% BSA for 2 hours at RT. cells were then washed and incubated with Alexa flour conjugated secondary antibodies (Alexa Fluor 488, Alexa Fluor 568, Alexa Fluor 647 conjugated) in PBS containing 2% BSA for 1 h at RT. After washing with PBS, cells were incubated with Dapi (1:10 000) for 15 min and after several washing steps mounted with Citifluor and visualize the staining pattern using a Leica fluorescence microscope.

### Hydroxyproline assays

The inferior lobe of right lungs was utilized to measure the collagen content using the hydroxyproline colorimetric assay kit according to the manufacturer instructions (Sigma, Catalog no. MAK008) as previously described (79). Briefly, the perfused lung lobes were weighed and hydrolyzed in 12N HCl for 16 hours at 120°C. Then, the hydrolysates of lung samples were incubated with chloramine T/oxidation buffer mixture followed by the addition of p–dimethylaminobenzaldehyde (DMAB) reagent. The hydroxyproline content was analysed by evaluating the absorbance values at 560 nm using a microplate reader. The quantity of hydroxyproline present in the lung tissue was determined as microgram (μg) of hydroxyproline per gram (g) lung tissue calculated through hydroxyproline standard curve.

### Enzyme-linked immunosorbent assay (ELISA)

BAL fluids collected from saline or bleomycin treated animals were assessed to measure the levels of cytokines IL-1β, IL-6 and TNFα using Mouse ELISA MAX™ Deluxe Set (Biolegend, catalog no. 432604, 431304 and 430904) according to the manufacturer protocol. The absorbance values were taken using a microplate reader and the concentration of the mentioned cytokines were calculated from their standard curve.

### Protein extraction and Western blot

The lung tissues and cultured cells from mice were homogenized in RIPA lysis buffer (Thermo Scientific, catalog no. 89901) supplemented with 1:100 ratio of protease and phosphatase inhibitor cocktail (Sigma, catalog no. 11836170001 and 4906845001). Protein lysates were then centrifuged at 13000 rpm for 15 minutes at 4°C, collected the supernatants and stored at −80°C until use. Total protein concentration was measured with the Pierce BCA protein assay kit (Thermo Scientific, catalog no. 23225) according to the manufacturer’s instructions. About 20-30 μg of total protein samples were placed on 10% SDS-polyacrylamide gel and transferred to nitrocellulose membrane using the Trans-Blot Turbo system (Bio-Rad). Membranes were incubated with 2-5% BSA blocking buffer in tris-buffered saline containing 0.1% Tween (TBST) and then incubated with primary antibodies diluted in TBST containing 2-5% BSA for overnight at 4°C. At next, blotting membranes followed washing thoroughly with TBST buffer and subsequently probed with the applicable horseradish peroxidase-conjugated secondary antibodies (Santa Cruz) in TBST containing 2% BSA for 1.5 hours at RT. Immunoreactive bands were then visualized by chemiluminescence reagents (Hiss GmbH, 16026) using Gel Doc XR+ System (Bio-Rad).

### RNA isolation, cDNA synthesis, and qRT-PCR

Total RNA from the mouse lung was isolated using TRIzol (Life Technologies, catalog no. 15596018) and total RNA from cultured cells was isolated using FavorPrep™ Tissue Total RNA Mini Kit (Favorgen Biotech, catalog no. FATRK 001-2). The concentration and quality of the RNA were assessed with UV spectrophotometry (NanoDrop Technologies, Wilmington, NC, USA). For cDNA synthesis, total RNA was reverse transcribed using random hexamers and SuperScript III FirstStrand Synthesis system (Life Technologies, catalog no. 18080-051). Gene expression experiment was conducted through quantitative RT-PCR (ABI PRISM 7900) using 10-μl reaction mixture containing 20 ng cDNA, SYBR Green Master Mix (Life Technologies, catalog no. 4368702), 10 μM forward primer, and 10 μM reverse primer. Results were analysed with Real-Time PCR System Software (Applied Biosystems). Gene expression data were normalized with the reference gene *glyceraldehyde-3 phosphate dehydrogenase* (*Gapdh*) or *β-actin* using the 2^−ΔΔCt^ method to determine the fold change.

### Flow cytometry

The lungs from mice were perfused and excised into very small pieces and subsequently digested with 600 U/mL collagenase II (Thermo Fischer, catalogue no. 17101015), DNase I 60 U/mL (Thermo Fischer, catalogue no. EN0521) in RPMI media (Lonza, Catalog no. 12-702F) at 37°C shaker with 200-220 rpm for 30-35 minutes. The cell suspension was then filtered through a 40-lM cell strainer (Sigma, catalogue no. CLS431750) and subsequently centrifuged. After washing the cell pellet with PBS containing 4% FBS, cells were blocked with FcR blocking reagent (Miltenyi, catalogue no. 130-092-575) for 10 min in ice. Cells were then incubated with a mixture of antibodies, PerCP/Cy5.5-conjugated CD45 (1:100), BV 650-conjugated CD326 (1:100), BV 711-conjugated CD64 (1:100), PE-conjugated CD31 (1:100), and eFluor 660-conjugated podoplanin (1:100) to sort the lung epithelial cells. Flow cytometric analyses were performed using a FACSAria II cell sorter (BD Bioscience, Singapore).

### Statistical Analysis

All experimental data are presented as mean ± standard error mean (SEM). Results were analyzed with unpaired t-test within two groups. Comparisons among three or more groups were calculated using one-way ANOVA, followed by Tukey’s post-hoc test. Differences were considered significant when the p-value was < 0.05. (*, *p<0.05*; **, *p<0.01*; ***, *p<0.001*; *NS*, not significant). Statistical analyses were performed using Graph-Pad Prism v.5 (GraphPad Software, USA).

### Antibodies

The following antibodies were used for immunostaining and western blot and flow cytometry:

| Target protein | Application | Source |
| --- | --- | --- |
| F4/80 | Immunostaining | Santa Cruz, catalog no. sc 377009 |
| YAP1 | Immunostaining | Proteintech, catalog no. 13584-1-AP |
| TAZ | Immunostaining, Western blot | Proteintech, catalog no. 23306-1-AP |
| Pro-SPC | Immunostaining | Merck, catalog no. AB3786 |
| Collagen I | Western blot | Affinity Biosciences, catalog no. AF7001 |
| Collagen I | Immunostaining | Novus Biologicals, catalog no. NB600-408 |
| IGF1R | Immunostaining, Western blot | Proteintech, catalog no. 20254-1-AP |
| MMP1 | Western blot | Proteintech, catalog no. 10371-2-AP |
| MMP9 | Western blot | Proteintech, catalog no. 10375-2-AP |
| MMP3 | Western blot | Proteintech, catalog no. 66338-1-Ig |
| TIMP1 | Western blot | Proteintech, catalog no. 30869-1-AP |
| TIMP2 | Western blot | Proteintech, catalog no. 17353-1-AP |
| CTGF | Western blot | Santa Cruz, catalog no. sc-365970 |
| PDGFRα | Western blot | Santa Cruz, catalog no. sc-398206 |
| Ki67 | Immunostaining | Cell Signaling, catalog no. 9449 |
| pAP-1 | Western blot | Proteintech, catalog no. 24909-1-AP |
| pSTAT3 | Western blot | Cell Signaling, catalog no. 9145 |
| pJNK | Western blot | Cell Signaling, catalog no. 4668 |
| pP50 | Western blot | Invitrogen, catalog no. 701430 |
| pP65 | Western blot | Cell Signaling, catalog no. 3033 |
| Vimentin | Immunostaining | Cell Signaling, catalog no. 5741 |
| DDR2 | Immunostaining | Santa Cruz, catalog no. sc-81707 |
| P16 | Immunostaining | Proteintech, catalog no. 10883-1-AP |
| P21 | Immunostaining | Proteintech, catalog no. 28248-1-AP |
| β-actin | Western blot | Santa Cruz, catalog no. sc-47778 |
| αSMA | Immunostaining, Western blot | Sigma, catalog no. A2547 |
| IGF1 | Western blot | Proteintech, catalog no. 28530-1-AP |
| CD45 PerCP/Cy5.5 | Flow cytometry | eBioscience, catalog no. 45-0451-82 |
| CD326 BV650 | Flow cytometry | BD Bioscience, catalog no. 740559 |
| Podoplanine Fluor<sup>TM</sup>660 | Flow cytometry | eBioscience, catalog no. 50-5381-82 |
| CD31 PE | Flow cytometry | Miltenyi Biotec, catalog no. 130-102-608 |

## Author Contributions

M.M.M., U.N. and A.S. conducted experiments and analyzed data. M.M.M. designed the experiments, oversaw the project, and wrote the initial draft of the manuscript. M.K.S. conceived and supervised the project, designed and interpreted all experiments, and wrote the final manuscript. All authors contributed to discussing the results and implications and provided feedback on the manuscript throughout its development.

## Acknowledgments

We express our gratitude to the members of the M.K.S. lab for their valuable discussions and insights. This work was supported by research funding from the Singapore Ministry of Health (MOH-OFIRG24jan-0018, MOH-001625), the Goh Foundation, and the Khoo Bridge Funding Award, awarded to MKS.

## Conflict of interest

The authors declare that there are no conflicts of interest.

---

**Supplementary Figure 1:**
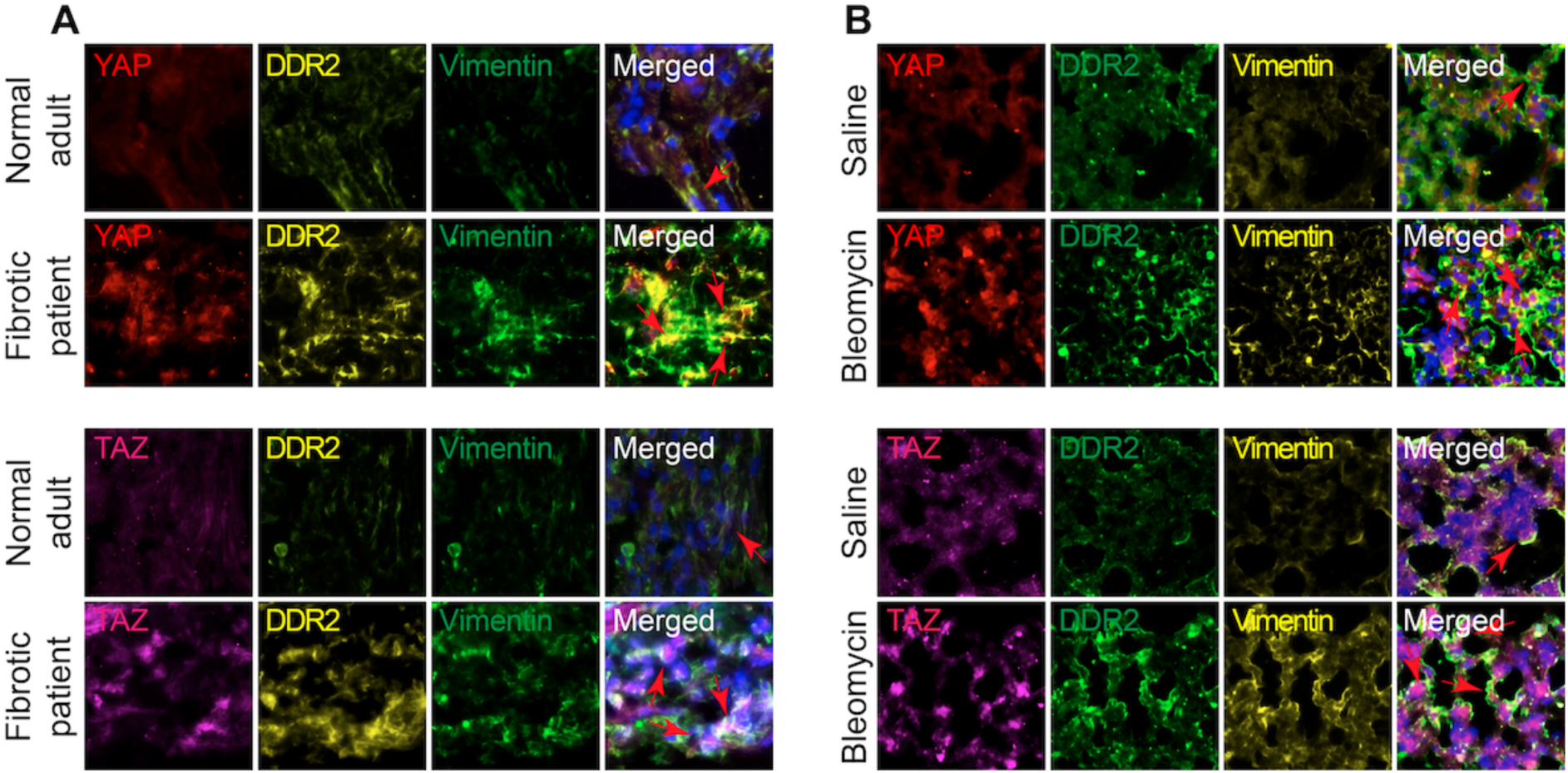
YAP/TAZ are activated lung fibroblasts in fibrosis. Representative immunofluorescence images from lung tissue sections after bleomycin-induced injury stained for DDR2 (yellow), Vimentin (green), YAP (red), TAZ (pink), and Dapi (blue). (A) Human patients diagnosed with pulmonary fibrosis showed increased nuclear presence of YAP or TAZ in lung fibroblasts compared with normal adult human lung. Red arrows indicate the cells that are positive for DDR2 and Vimentin together with YAP or TAZ. (B) Lung fibroblasts showed increased nuclear presence of YAP or TAZ at 7 days after bleomycin treatment. Red arrows indicate the cells that are positive for DDR2 and Vimentin together with YAP or TAZ in bleomycin-induced fibrotic lung.

**Supplementary Figure 2:**
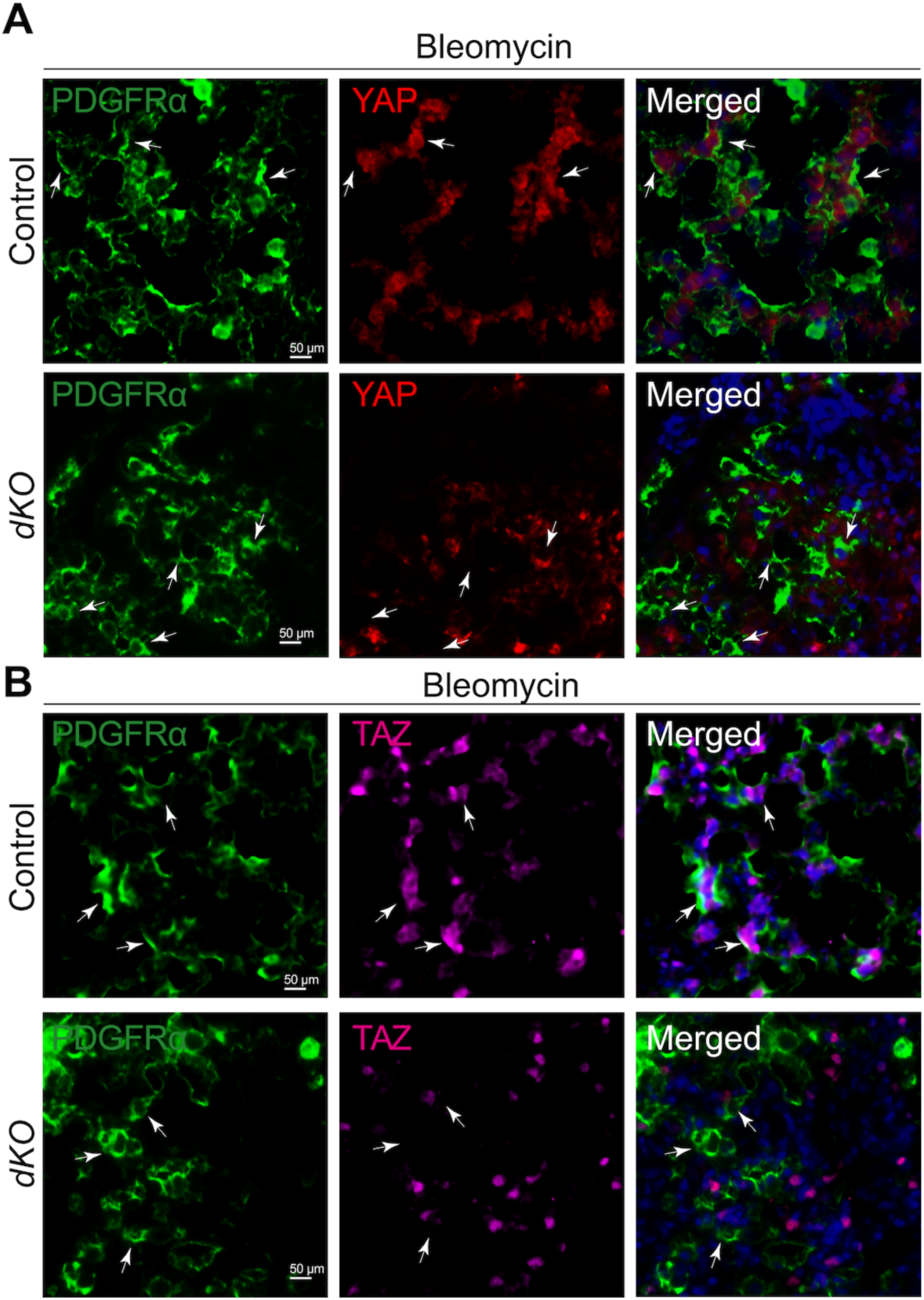
Bleomycin treatment activates the YAP/TAZ expression in lung fibroblasts. (A-B) Representative images from lung tissue sections after bleomycin-induced injury stained for PDGFRα (green), YAP (red), TAZ (pink), and Dapi (blue). Lung fibroblasts showed increased nuclear presence of YAP and TAZ at 7 days after bleomycin treatment in control mice. In *dKO* lung, the expression of YAP and TAZ were inactivated in PDGFRα fibroblasts. White arrows indicate the cells that are positive for PDGFRα together with YAP or TAZ in controls lung section; however, in *dKO* lung white arrows indicate the cells that are positive for PDGFRα only, but negative for YAP or TAZ expression.

**Supplementary Figure 3:**
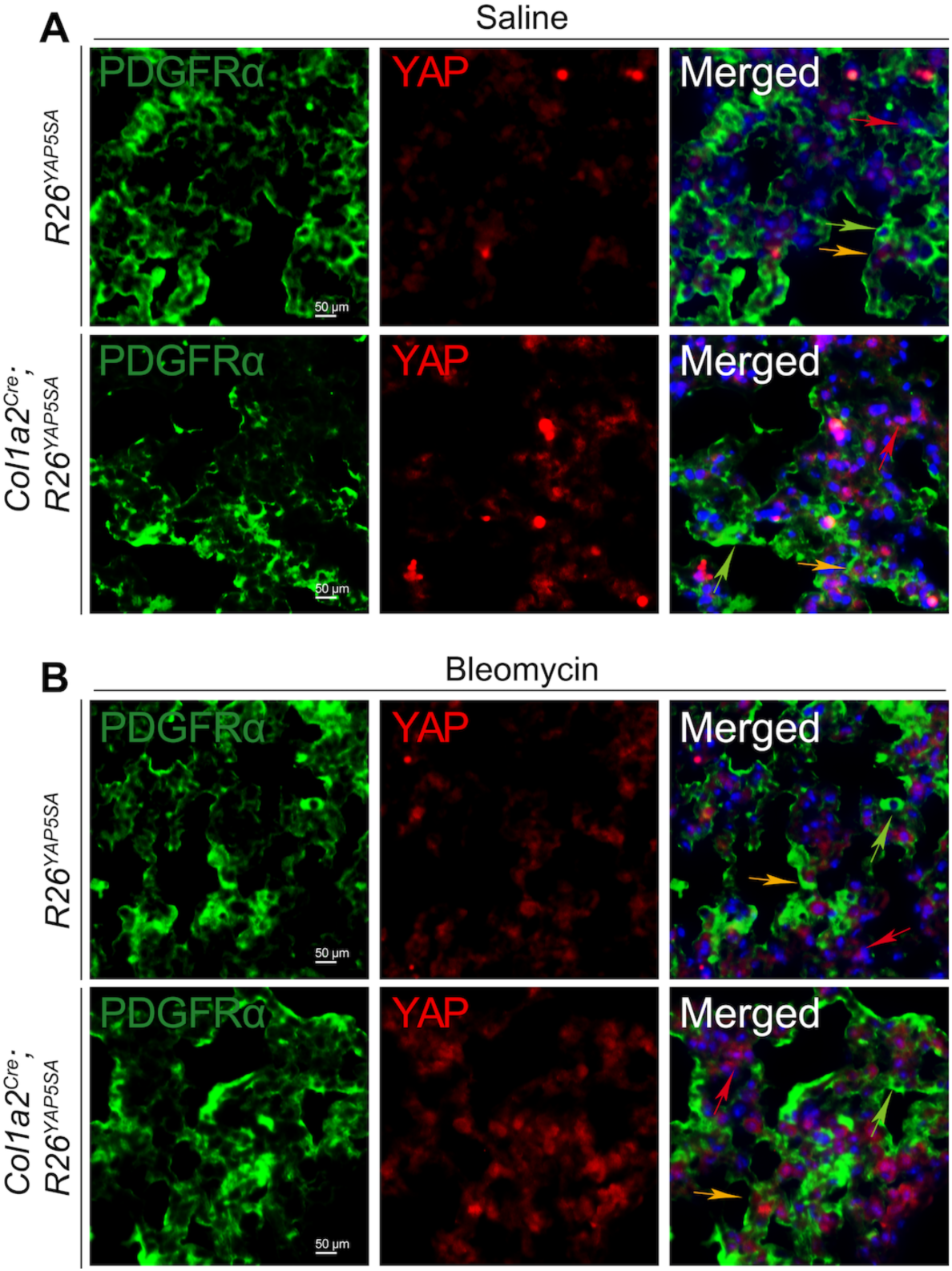
YAP is hyperactivated in lung fibroblasts after bleomycin treatment in *Col1a2^Cre(ER)T^; R26^YAP5SA^* mice. Representative images from lung tissue sections after saline or bleomycin-induced injury stained for PDGFRα (green), YAP (red), and Dapi (blue). In comparison to *R26^YAP5SA^* lungs, *Col1a2^Cre(ER)T^; R26^YAP5SA^* lungs showed increased nuclear localization of YAP at 14 days after saline or bleomycin treatment; although the nuclear localization of YAP was more prominent after bleomycin-induced injury. Green arrows indicate the cells that are positive for PDGFRα only; red arrows indicate the cells that are positive for YAP only; yellow arrows indicate the cells that are positive for PDGFRα, YAP, and Dapi.

**Supplementary Figure 4:**
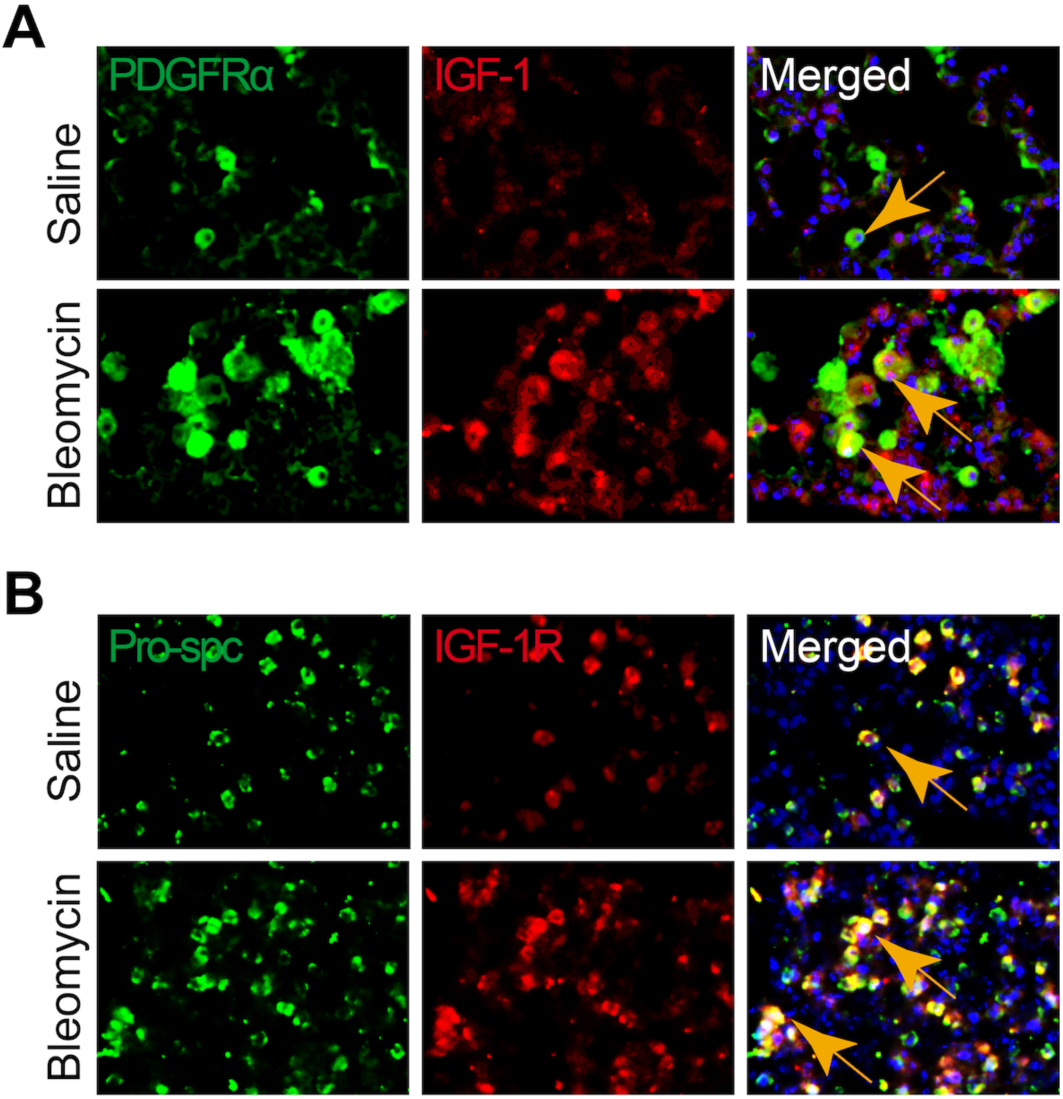
Expression of IGF1 in fibroblasts and IGF1R in AT2 cells after bleomycin-injury. (A) Immunofluorescence staining of IGF1 co-stained with PDGFRα in wildtype mice lung sections at 14 days post-bleomycin injury compared to saline-treated controls (n = 5). (B) Immunofluorescence staining of IGF1R co-stained with Pro-SPC in wildtype mice lung sections at 14 days post-bleomycin injury compared to saline-treated controls (n = 5).

**Supplementary Figure 5:**
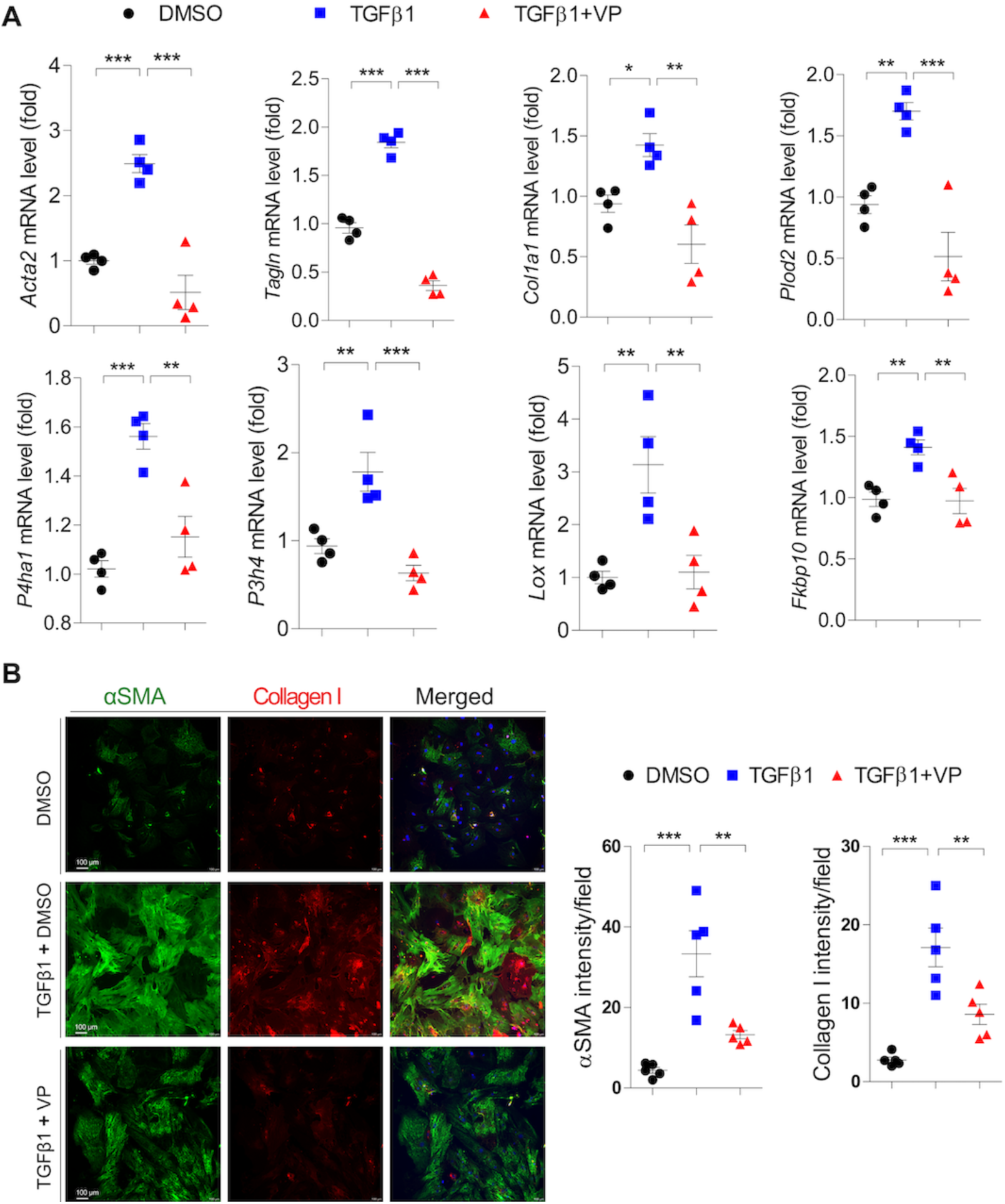
Verteporfin impairs myofibroblasts formation and collagen I production. (A) Real-time qPCR of *Col1a1*, *Acta2*, *SM22a*, *Plod2*, *P3h4*, *Lox* and *Fkbp10* on primary lung fibroblasts treated with/without TGFβ1 alone or TGFβ1+VP condition for 96 hours. (B) Immunofluorescence staining and quantification of αSMA and collagen I using primary mouse lung fibroblasts treated for 96 hours with TGFβ1 or TGFβ1+VP (co-treatment). DMSO was used as vehicle control. Fluorescence intensity was quantified with imageJ software. The data are represented as the mean ± SEM; comparison by one-way ANOVA, followed by Tukey’s post-hoc test. *, *p* < 0.05; **, *p* < 0.01; ***, *p* < 0.001; NS, not significant.

**Supplementary Figure 6:**
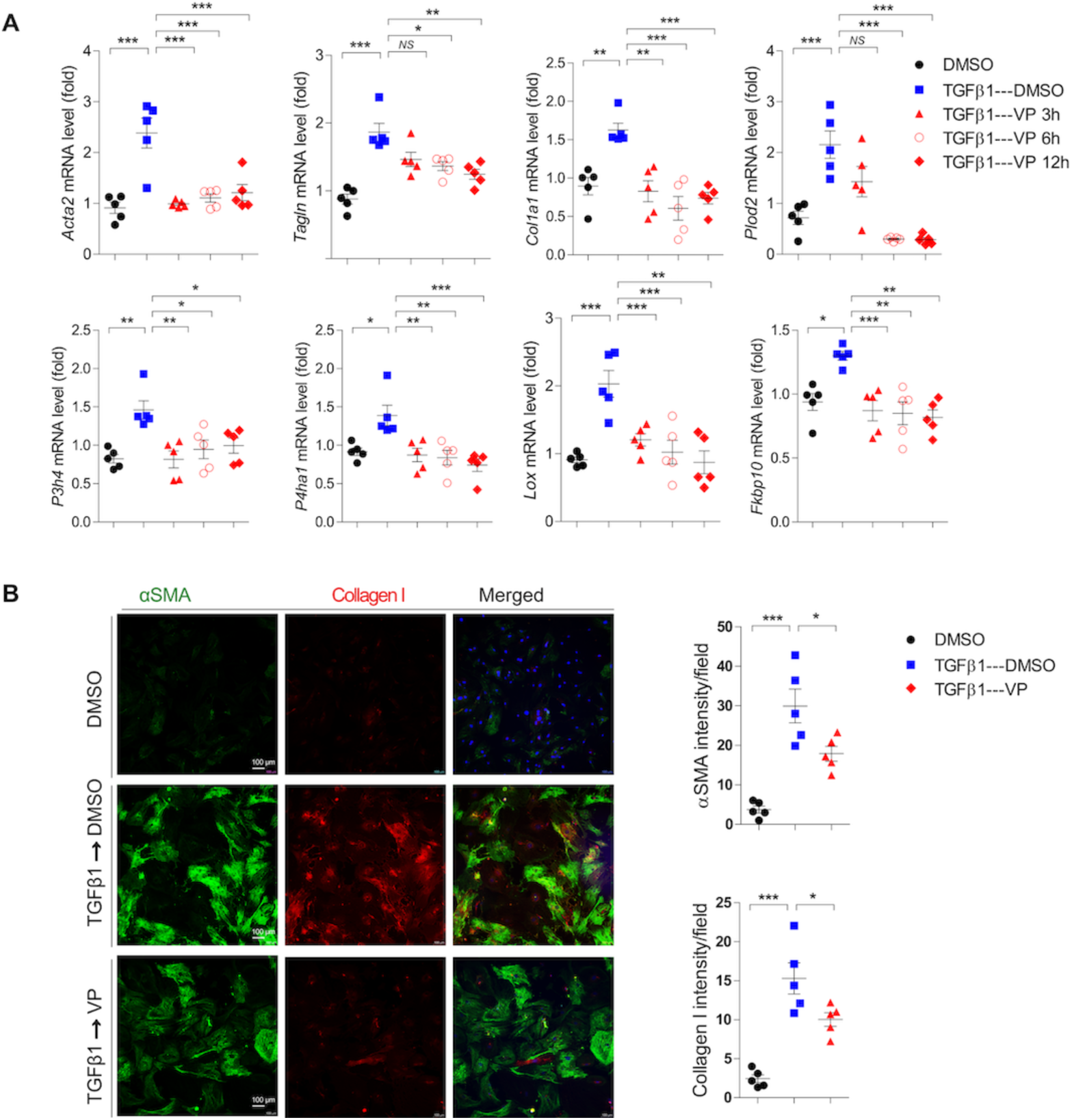
Verteporfin impairs myofibroblasts formation and collagen I production. (A) Real-time qPCR of *Col1a1*, *Acta2*, *SM22a*, *Plod2*, *P3h4*, *Lox* and *Fkbp10* on primary lung fibroblasts cultured for 96 hours in the presence of TGFβ1, followed by a post-treatment with VP or DMSO as indicated. (B) Immunofluorescence staining and quantification of αSMA and collagen I using primary mouse lung fibroblasts cultured for 96 hours in the presence of TGFβ1, followed by a post-treatment with VP or DMSO for 24 hours. Fluorescence intensity was quantified with imageJ software. The data are represented as the mean ± SEM; comparison by one-way ANOVA, followed by Tukey’s post-hoc test. *, *p* < 0.05; **, *p* < 0.01; ***, *p* < 0.001; NS, not significant.

**Supplementary Figure 7:**
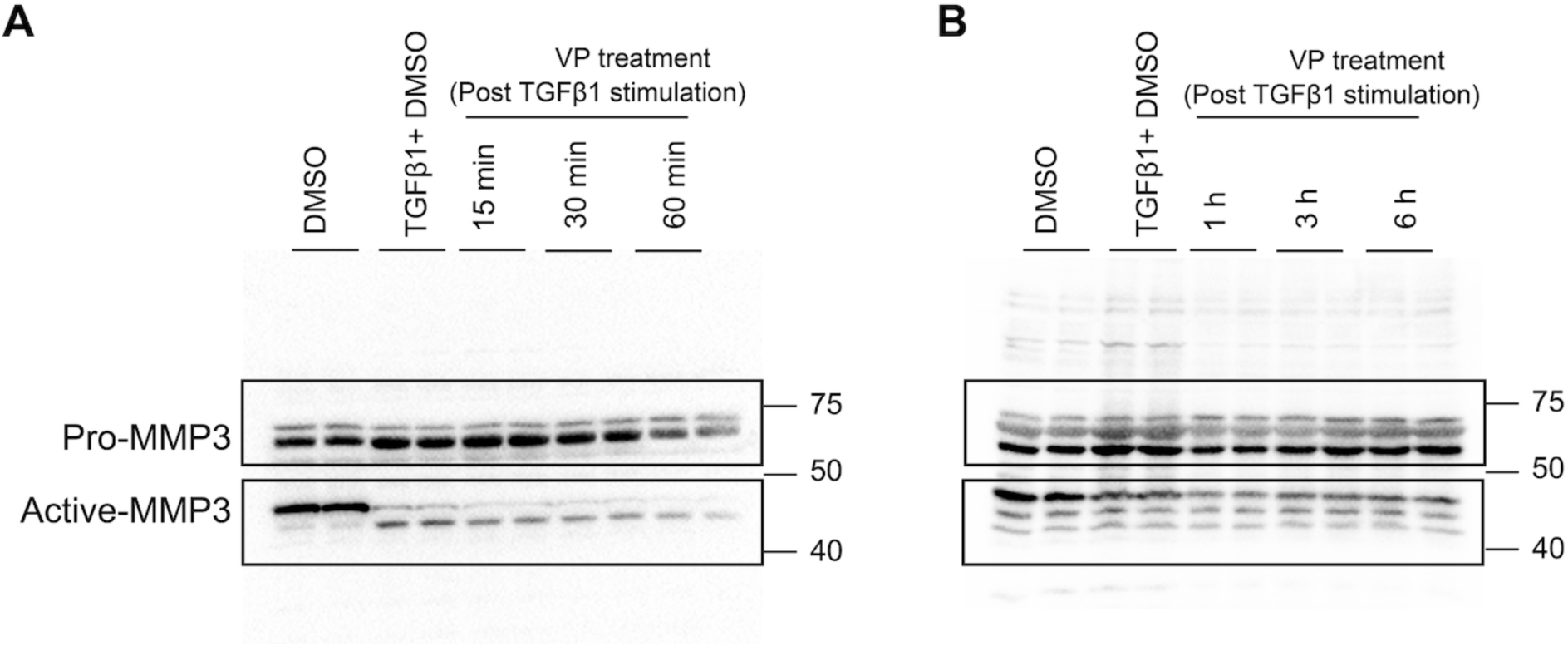
Verteporfin attenuates the active form of MMP3 in myofibroblasts. Primary mouse lung fibroblasts were cultured for 96 hours in the presence of TGFβ1, followed by a post-treatment with VP for 15, 30, 45 and 60 minutes respectively. In a separate experiment, post-treatment with VP in the presence of TGFβ1 was also performed for 1, 3 and 6 hours respectively. DMSO was used as vehicle control. Immunoblot analysis of MMP3 protein was performed on the indicated experiments.

## Notes

### Competing Interest Statement

The authors have declared no competing interest.

